# Genomic underpinnings of convergent adaptation to high altitudes for alpine plants

**DOI:** 10.1101/2022.10.20.508685

**Authors:** Xu Zhang, Tianhui Kuang, Wenlin Dong, Zhihao Qian, Huajie Zhang, Jacob B. Landis, Tao Feng, Lijuan Li, Yanxia Sun, Jinling Huang, Tao Deng, Hengchang Wang, Hang Sun

**Affiliations:** CAS Key Laboratory of Plant Germplasm Enhancement and Specialty Agriculture, Chinese Academy of Sciences, Wuhan Botanical Garden, Wuhan, Hubei 430074, China; Center of Conservation Biology, Core Botanical Gardens, Chinese Academy of Sciences, Wuhan, Hubei 430074, China; Yunnan International Joint Laboratory for Biodiversity of Central Asia, Key Laboratory for Plant Diversity and Biogeography of East Asia, Kunming Institute of Botany, Chinese Academy of Sciences, Kunming, Yunnan 650201, China; School of Integrative Plant Science, Section of Plant Biology and the L. H. Bailey Hortorium, Cornell University, Ithaca, NY 14850, USA; BTI Computational Biology Center, Boyce Thompson Institute, Ithaca, NY 14853, USA; State Key Laboratory of Crop Stress Adaptation and Improvement, School of Life Sciences, Henan University, Kaifeng, Henan 475004, China; Department of Biology, East Carolina University, Greenville, NC 27858, USA; University of Chinese Academy of Sciences, Beijing 100049, China

**Keywords:** alpine plants, convergent adaptation, evolutionary rates, genomics, “greenhouse” morphology, molecular convergence

## Abstract

Evolutionary convergence is one of the most striking examples of adaptation driven by natural selection. However, genomic evidence for convergent adaptation to extreme environments remains scarce. The Himalaya-Hengduan Mountains represent the world’s most species-rich temperate alpine biota, providing an ideal “natural laboratory” for studying convergent adaptation to high altitudes. Here, we generate reference genomes for two alpine plants, *Saussurea obvallata* (Asteraceae) and *Rheum alexandrae* (Polygonaceae), with 37,938 and 61,463 annotated protein-coding genes. By integrating an additional five alpine genomes, we investigate genomic signatures of convergent adaptation to the hostile environments of high altitudes. We show that alpine genomes tend to mitigate their genetic load by contracting genes functioning in the immune system to survive such harsh environments with few pathogens present. We detect signatures of convergent positive selection on a set of genes involved in reproduction and development and reveal that molecular convergence has acted on genes involved in self-incompatibility, cell wall modification, DNA repair and stress resistance, which underlie adaptation to extremely cold, high UV radiation and hypoxia environments. Using gene expression profiles, we further demonstrate that genes associated with cuticular wax and flavonoid biosynthetic pathways exhibit higher expression levels in leafy bracts, shedding lights on the genetic mechanisms of the adaptive ‘greenhouse’ morphology. Our integrative data provide genomic insights into the convergent evolution at higher-taxonomic levels, aiding in deep understanding of genetic adaptation to complex environments.

## Introduction

Evolutionary biologists have long aimed to understand the extent that evolutionary trajectories are predictable, i.e., the extent to which convergent adaptation in distinct lineages is driven by conserved molecular changes (Zhang and Kumar 1997; Stern and Orgogozo 2009; Zhen et al. 2012; Storz 2016). Evolutionary convergence in settings where different species repeatedly face common selective pressures offer a powerful opportunity to address this issue (Yeaman et al. 2016; Birkeland et al. 2020; Xu et al. 2020). An ideal system for investigating the genetic underpinnings of convergent evolution is the independent adaptation of divergent lineages to high-altitude environments.

The Himalaya-Hengduan Mountains (HHMs) exhibit extraordinarily high species richness, and are believed to be the center and origin of diversity of many organisms in the Northern Hemisphere (Spicer et al. 2020). In the alpine zones of the HHMs (elevation above 4500 m), usually characterized by freezing temperatures, high UV (ultra violet) radiation, and hypoxia, plants typically possess suites of similar morphological and physiological adaptations to allow them to survive and reproduce in the hostile environments (Tsukaya and Tsuge 2001). In comparison to plants of lowland areas, plants living in the HHMs (and other high-altitude areas) have dwarf stems, smaller leaves and higher densities of branches, and often exhibit a specialized morphology such as leafy bracts, woolly coverings and cushion forms (Nagy and Grabherr 2009; Sun et al. 2014). Genome-wide studies have documented some genomic footprints of high-altitude adaptation by testing for positive selection and mining expended gene families, often involving functional pathways such as DNA repair, abiotic stress response, reproductive processes, as well as secondary metabolite biosynthesis (Zeng et al. 2015; Zhang et al. 2019; Wang et al. 2021). However, the limited availability of reference genomes for alpine plants restricts further understanding in the genomic evolution of high-altitude adaptation. Additionally, the genomic convergence underpinning high-altitude adaptation have not been examined.

To gain genomic insights into convergent adaptation to high-altitude environments, we newly assembled and annotated reference genomes of *Saussurea obvallata* (Asteraceae) and *Rheum alexandrae* (Polygonaceae). These two species are mainly found in mountain slopes and alpine meadows of the HHMs and are renowned for their ‘glasshouse’ morphology, i.e., the upper leaves of which have developed into large semi-translucent leafy bracts that cover the inflorescences, which have been shown to have significant ecological benefits to the plant (Song et al. 2015). We integrate an additional five available genomes of alpine plants [*Crucihimalaya himalaica* (Brassicaceae), *Eutrema heterophyllum* (Brassicaceae), *Hordeum vulgare* var. *nudum* (Poaceae), *Prunus mira* (Rosaceae) and *Salix brachista* (Salicaceae)] as well as their lowland relatives for comparative genomic analyses (supplementary table S1). These seven alpine species represent major clades of angiosperms that independently colonized high-altitude environments. Different from previous case studies of plant genomes, we take advantage of a comprehensive genomic data set of alpine plants to characterize genome-wide signatures of convergent evolution. Specifically, we intended to address three main questions: (i) Whether expanded or contracted gene families show a convergent pattern in alpine plants and have effects on the high-altitude adaptation? (ii) Which genes have undergone convergent molecular evolution and are involved in adaptation to the extremely cold, high UV radiation and hypoxia environment? (iii) Lastly, what are the genomic bases underlying the adaptive “greenhouse” morphology? In addressing these questions, this study provides novel insights into the genomic convergence that contribute to the adaptation to extreme environments.

## Results and Discussion

### Assembly and annotation of two reference genomes of alpine plants

Using a *K*-mer analysis method, we first estimated the genome size of *S. obvallata* and *R. alexandrae* to be ∼2,251 Mb and 2,137 Mb, respectively (supplementary figs. S1, S2, supplementary table S2). Then we obtained a chromosome-level genome of *S. obvallata* and a contig-level genome of *R. alexandrae* using Illumina, Oxford Nanopore, and high-throughput chromatin conformation contact (Hi-C) sequencing technologies (supplementary table S3). For the *S. obvallata* genome, a total of ∼95 Gb Illumina short reads, ∼143 Gb Nanopore long reads and ∼206 Gb Hi-C data were obtained. For *R. alexandrae* genome, a total of ∼104 Gb Illumina short reads and ∼144 Gb Nanopore long reads were generated. Additionally, transcriptomic data of *R. alexandrae* (∼25 Gb) and *S. obvallata* (∼157 Gb) were obtained for transcript-based gene annotation (supplementary table S3). Tissue-specific transcriptomic data of *S. obvallata* were also used for further gene expression analysis (supplementary table S4).

A *de novo* assembly pipeline allowed us to achieve initial genome assemblies that captured 2,044 Mb and 2,040 Mb in 145 and 129 contigs for *S. obvallata* and *R. alexandrae* genomes, with contig N50 of 36.96 Mb and 36.32 Mb, respectively (table 1; supplementary table S5). Using a Hi-C assisted assembly pipeline, 1,952 Mb which accounted for 95.5% of the assembled *S. obvallata* genome, was anchored on 16 chromosomes (table 1; supplementary figs. S3, S6), in line with previous cytological evidence (Fujikawa et al. 2004). We further evaluated the completeness of the assembled genomes and found high completeness rates (94.6% of *S. obvallata* and 94.1% of *R. alexandrae*) of both assemblies as evidenced by BUSCO (Benchmarking Universal Single-Copy Orthologs) assessments using the Eukaryota_odb10 database (table 1; supplementary tables S7, S8) (Manni et al. 2021). The long terminal repeat (LTR) assembly index (LAI), which evaluates the contiguity of intergenic and repetitive regions of genome assemblies based on the intactness of LTR retrotransposons (Ou et al. 2018), was 19.68 for *S. obvallata* and was 15.01 for *R. alexandrae*, respectively, showing high continuity of both genomes.

**Table 1.** Statistics of genome assembly and annontation of *S. obvallata* and *R. alexandrae*.

| Statistic | *S. obvallata* | *R. alexandrae* |
| --- | --- | --- |
| **Total length (bp)** | 2,044,030,733 | 2,039,881,226 |
| **Number of contigs** | 145 | 129 |
| **Largest contig (bp)** | 126,457,859 | 160,815,976 |
| **Anchored length (bp)** | 1,951,503,694 | - |
| **GC (%)** | 37.94 | 41.41 |
| **N50 (bp)** | 36,958,263 | 36,323,674 |
| **Complete BUSCOs (%)** | 94.6 | 94.1 |
| **Repeat content (%)** | 81.88 | 81.65 |
| **Number of genes** | 37,938 | 61,463 |

Transposable elements (TEs) and other repeat sequences accounted for 81.88% and 81.65% of the *S. obvallata* and *R. alexandrae* assemblies, respectively (table 1). In *S. obvallata*, LTR retrotransposons (43.95%), followed by DNA TEs (2.03%) and LINEs (0.93%), were most abundant, with LTR-*Gypsy* and LTR-*Copia* retrotransposons accounting for 18.19% and 25.76% of the LTRs, respectively (supplementary table S9). LTR retrotransposons (44.31%), LINES (2.73%), and DNA TEs (4.08%) accounted for most of the *R. alexandrae* repeats, with LTR-*Gypsy* (37.98%) and LTR-*Copia* (6.33%) retrotransposons predominant among the LTRs (supplementary table S10). A combination of transcript-based, *de novo* and homology-based prediction methods yielded 37,938 and 61,463 high-confidence protein-coding gene models (table 1; supplementary figs. S4, S5). By comparing to public protein databases, a total of 36,542 (96.32%) and 47,535 (77.34%) predicted genes of *S. obvallata* and *R. alexandrae* were functionally annotated (supplementary table S11). For the annotated genes, 93.5% and 96.1% of the complete BUSCO genes against the Eukaryota_odb10 database could be identified in *S. obvallata* and *R. alexandrae*, respectively. Overall, our newly assembled and annotated genomes of *S. obvallata* and *R. alexandrae* were of high quality, providing valuable genomic resources for the understanding of the convergent adaptation of alpine plants to high-altitude environments.

### Complex adaptive histories of alpine plants

The high-quality genomes of *S. obvallata* and *R. alexandrae* allowed reconstruction of the phylogenomic relationships and the adaptive histories of alpine lineages. We downloaded annotated protein sequences from genomes of five additional alpine species as well as 13 representative sisters living in low elevations (supplementary table S1). Species were phylogenetically widely dispersed with 17 eudicots and three monocots, placed in seven families of angiosperms. Orthologs were inferred using OrthoFinder (supplementary table S12), resulting in 6,711 gene families present in all 20 species, of which 195 single-copy orthogroups were used for phylogeny inference and divergence time estimation. A robustly supported phylogenetic tree obtained through maximum-likelihood (ML) analysis of the concatenated protein sequences was consistent with the known phylogenetic relationships within angiosperms, in which the alpine taxa included in our study occurred in seven independent lineages placed in six families (fig. 1a). The time tree inferred from MCMCTree showed a wide divergence history between alpine species and their sampled sisters, ranging from 2 million years ago (Mya) to 31.81 Mya (fig. 1a). This indicted multiple times of adaptation during a wide range of geological history for alpine plants to the high-altitude environments of the HHMs.

**FIG. 1.**
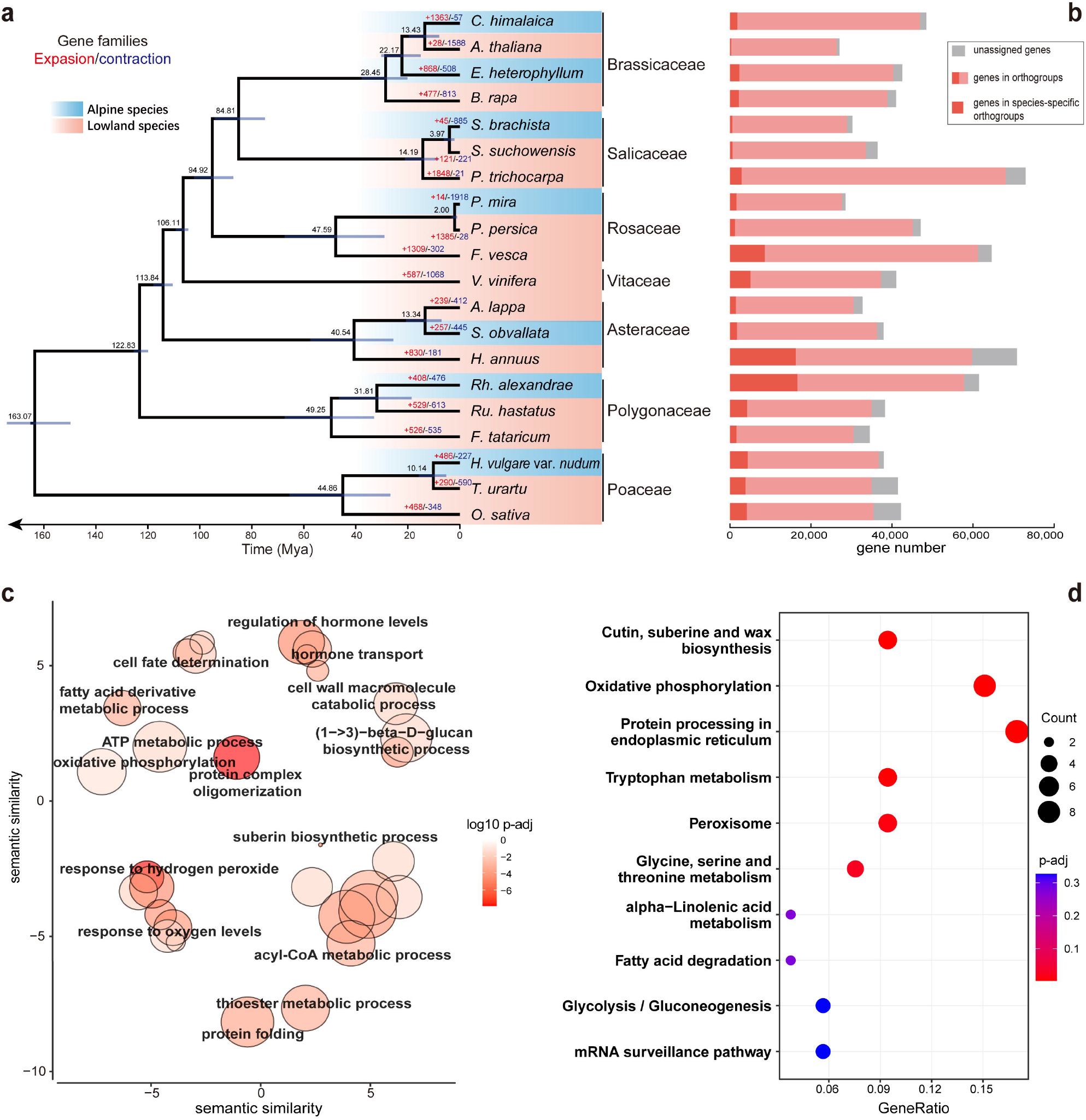
Evolutionary history of alpine plants. (a) Chronogram showing divergence times among alpine plants (cyan background) with their lowland relatives (orange background) with node age and 95% confidence intervals (blue bars). The red and blue numbers above the branches represent significant expansion and contraction events, respectively. (b) Bar plot showing gene number identified by OrthorFinder. (c) REVIGO clusters of significantly enriched GO terms for convergently expended (CoEx) gene families in alpine plant genomes. Each bubble represents a summarized GO term from the full GO list by reducing functional redundancies, and their closeness on the plot reflects their closeness in the GO graph, i.e., the semantic similarity. (d) KEGG pathways of CoEx gene families in alpine plant genomes. “p-adj” refers to the adjusted *p*-value using the Benjamini– Hochberg method.

### Convergent changes of gene family number

We determined convergent changes in gene family number when a gene family showed significant expansion or contraction in more than three alpine species. A total of 56 convergently expanded (CoEx) gene families were identified. Biological Process (BP) of Gene Ontology (GO) and Kyoto Encyclopedia of Genes and Genomes (KEGG) analyses of CoEx families found 68 significantly enriched GO terms and seven KEGG pathways (*p*-adjust < 0.05) (supplementary tables S13, S14). Enriched pathways of CoEx gene families were mainly related to abiotic resistance, such as response to hypoxia, regulation of hormone levels and hormone transports (figs. 1c, 1d; supplementary figs. S6, S7). Interestingly, we found a greater number of convergently contracted (CoCo) families than CoEx families, with 1,193 gene families convergently contracted, involving 390 significantly enriched GO terms and 27 KEGG pathways (*p*-adjust < 0.05) (supplementary tables S15, S16). Most of these pathways included genes involved in the response to biotic stresses, such as defense to pathogens and toxicants (supplementary figs. S8, S9).

A unique stress at high altitudes is hypobaric hypoxia (Beall 2014). In plants, an oxygen deficiency dramatically reduces the efficiency of cellular ATP production, which has diverse ramifications for cellular metabolism and developmental processes (Fukao and Bailey-Serres 2004). Various oxygen-sensing mechanisms have been described that are thought to trigger plant response to low-level oxygen and thus adaptation to altitude (Abbas et al. 2022). In our analysis, we found many significantly enriched GO terms of CoEx families were functionally related to the response to oxygen levels, such as response to hypoxia, response to decreased oxygen level and response to hydrogen peroxide (fig. 1c). These pathways included genes encoding the alcohol dehydrogenase (ADH) (Peng et al. 2001), and the HSP20-like chaperones superfamily proteins that are functionally enriched in the GO term, cellular response to hypoxia. In *A. thaliana*, the oxygen-sensing system is mediated by the plant cysteine oxidase (PCO) N-degron pathway substrates group VII ethylene response factors (ERF-VIIs) (Licausi et al. 2011), which are involved in modulating ethylene response activating the expression of *ADH1* (Yang et al. 2011). While we did not discover convergent expansion of ERF-VIIs in alpine plants, we found genes that are significantly enriched GO terms related to response to ethylene, such as *ADH1, ERF1* (*ETHYLENE RESPONSE FACTOR 1*) and *EER1* (*ENHANCED ETHYLENE RESPONSE 1*). Nonetheless, the expansion of genes involved in the response to hypoxia is necessary for the adaptation of alpine plants to low-level oxygen in high-altitude environments.

In addition, multiple CoEx families were found to be significantly enriched in plant hormone pathways. Examples include genes encoding the probable indole-3-pyruvate monooxygenases (YUC) involved in auxin biosynthesis (Cao et al. 2019), and the small auxin up-regulated RNAs (SAUR) and the D6 protein kinase (D6PK) involved in auxin polar transport (van Berkel et al. 2013). Auxin regulates a series of developmental processes such as apical dominance, plant organogenesis, and reproductive development by affecting cell growth, differentiation, and patterning (Mockaitis and Estelle 2008). Other developmental regulation pathways including leaf senescence, phototropism and plant organ senescence were also detected to be significantly enriched in CoEx families. This result, coupled with the commonly observed morphological divergence between alpine plants and lowland relatives (Sun et al. 2014), indicates that regulation of morphological changes can be a pivotal path for plant adaptation to high-altitude environments.

Many CoCo families in alpine plants were found to be functionally related to response to pathogens or toxicants, involving GO terms of response to toxic substance and oomycetes, xenobiotic transport and detoxification (supplementary tables S15, S16). Related gene families include the cysteine-rich receptor-like kinases (CRKs), a large subfamily of receptor-like protein kinases (RLKs) that play vital roles in defense responses and programmed cell death in plants (Chen et al. 2004), the malectin-like receptor kinases (MLRs) orchestrating the plant immune responses and the accommodation of fungal and bacterial symbionts (Ortiz-Morea et al. 2022), the multidrug and toxic compound extrusion (MATE) proteins involved in xenobiotic detoxification and multidrug resistance (Diener et al. 2001), as well as the ubiquitin-conjugating (E2) enzymes emerging in recent years as an important regulatory factor underlying plant innate immunity (Zhou et al. 2017). Moreover, the results also showed that the genes encoding receptors for tyrosine-sulfated glycopeptide (PSY1Rs) and phytosulfokine (PSKRs), belonging to the leucine-rich repeat receptor kinases (LRR-RKs), have been undergoing contraction in all alpine species. PSKR1 and PSY1R have been shown to involve in plant immune, with antagonistic effects on bacterial and fungal resistances (Mosher et al. 2013).

The largest disease-resistance genes comprise genes encoding nucleotide-binding site and leucine-rich-repeat domain receptors (NBS-LRRs). Three NBS-LRR gene subclasses, TIR-NBS-LRR (TNL), CC-NBS-LRR (CNL), and RPW8-NBS-LRR (RNL), have been characterized based on the N-terminal domains (McHale et al. 2006). We manually annotated NBS-LRR genes in the sampled genomes using the HMMER search with the Pfam database (Wheeler and Eddy 2013). A total of 5,655 NBS-LRR genes, including 4,058 CNLs, 1,024 TNLs and 173 RNLs were identified among all analyzed genomes (supplementary fig. S10). Additionally, we reconciled the NBS-LRR gene tree to examine the gains and losses of NBS-LRR genes in alpine species. The results showed that, compared to lowland relatives, most alpine species tend to lose more NBS-LRR genes and exhibit reduced copy number, while *E. heterophyllum, S. obvallata* and *R. alexandrae* have similar numbers compared to their closest relatives (supplementary fig. S11). A similar phenomenon was also described in the case study of the *C. himalaica* genome, in which the most significantly contracted gene families were functionally enriched in disease and immune responses pathways (Zhang et al. 2019). Due to the harsh environments characterized by freezing temperatures, aridity, and high UV radiation, it is reasonable to hypothesize that gene families involved in pathogen or toxicant defense have undergone contraction in alpine plants, as fewer microorganisms exist. In addition, genes functioning in cellular transport were shown to be contracted in alpine plants. Related pathways include cytoskeleton organization, actin filament-based process, export across plasma membrane and export from the cell (fig. 1d; supplementary table S15). These processes may be a component of plant immune system and possibly have undergone simplification due to the pathogen depauperate environments of high altitudes. These results suggest that contractions immune system genes might be an effective way to mitigate genetic loads while surviving in hostile environments but with few pathogens present.

### Tests for convergent positive selection

In harsh environments, positive selection is expected to be common in genes controlling early life history stages, including genes involved in reproduction and development (Cui et al. 2019). Our branch-site tests identified 36 convergently selected genes that show signatures of positive selection in more than three alpine species (supplementary table S17). These genes were functionally related to basal life processes involving reproduction and respiration, such as carpel, gynoecium, ovule and endosperm developments and photorespiration pathways (supplementary fig. S12, supplementary tables S18, S19). Examples included the *SEPALLATA* (*SEP*) MADS-box genes required in floral organ and meristem identity (Pelaz et al. 2000), the *MALE MEIOCYTE DEATH 1* (*MMD1*) gene regulating cell cycle transitions during male meiosis (Yang et al. 2003), the *NIJMEGEN BREAKAGE SYNDROME 1* (*NBS1*) gene involved in double-strand break repair, DNA recombination and maintenance of telomere integrity in the early stages of meiosis (Zhang et al. 2006), and the *HYDROXYPYRUVATE REDUCTASE* (*HPR*) gene localized in leaf peroxisomes functioning in the glycolate pathway of photorespiration (Mano et al. 1999). Moreover, genes encoding pectin methylesterases (PMEs) were detected to be undergone convergently positive selection (supplementary table S17). PMEs play a central role in the synthesis and metabolism of pectins, which contribute both to the firming and softening of the cell wall that is important for basal developmental processes of higher plants, such as meristematic growth, fruit ripening, programmed cell death, and endosperm rupture upon germination (Louvet et al. 2006; Rodriguez-Gacio Mdel et al. 2012). In addition, the glyoxylate and dicarboxylate metabolism, the most significantly enriched KEGG pathways (supplementary fig. S13, supplementary table S19), is a fundamental biochemical process that ensures a constant supply of energy to living cells. These convergently selected genes detected in our analyses likely contribute to the primary adaptation of alpine plants to similar extreme environments.

### Detection of molecular convergence

We investigated signatures of genes undergoing molecular convergence among alpine species using a combination of approaches for the detection of convergent evolutionary rate shifts and site-based estimation of convergent amino acid (AA) evolution. These approaches have been commonly used in previous studies that investigate the genomic signatures of convergent evolution, including convergent adaptation to seasonal habitat desiccation in African killifishes (Cui et al. 2019), convergent regulatory evolution and loss of flight in paleognathous birds (Sackton et al. 2019), and convergent evolution of extreme lifespan in Pacific Ocean rockfishes (Kolora et al. 2021).

Convergent shifts in gene evolutionary rates were detected using the RERconverge method (Kowalczyk et al. 2019), which estimates the correlation between relative evolutionary rates (RERs) of protein sequences and the evolution of a convergent binary or continuous trait across a phylogeny. Our analysis focused on positive correlations, representing genes with faster evolutionary rates in alpine species relative to species living in low elevations. An increased RER could arise due to relaxation of a constraint or positive selection, which could be adaptive to habitat-related changes (Kowalczyk et al. 2020). We identified 69 gene families undergoing convergently accelerated RERs in alpine species, significantly enriched in 93 GO terms and 12 KEGG pathways (*p*-adjust < 0.05) (supplementary tables S20, S21), involving pathways for self-incompatibility system, cell wall modification, DNA repair and stress resistance (fig. 2). Among them, 26 gene families were found to have convergent AA shifts by a PCOC analysis (supplementary fig. S14). The PCOC method considers shifts in AA preference instead of identical substitutions (Rey et al. 2018). Given the relatively far phylogenetic distances among our analyzed species, selecting only sites that converged to the exact same AA in all species is quite strict and is bound to capture only a subset of the substitutions associated with the convergent trait change.

**FIG. 2.**
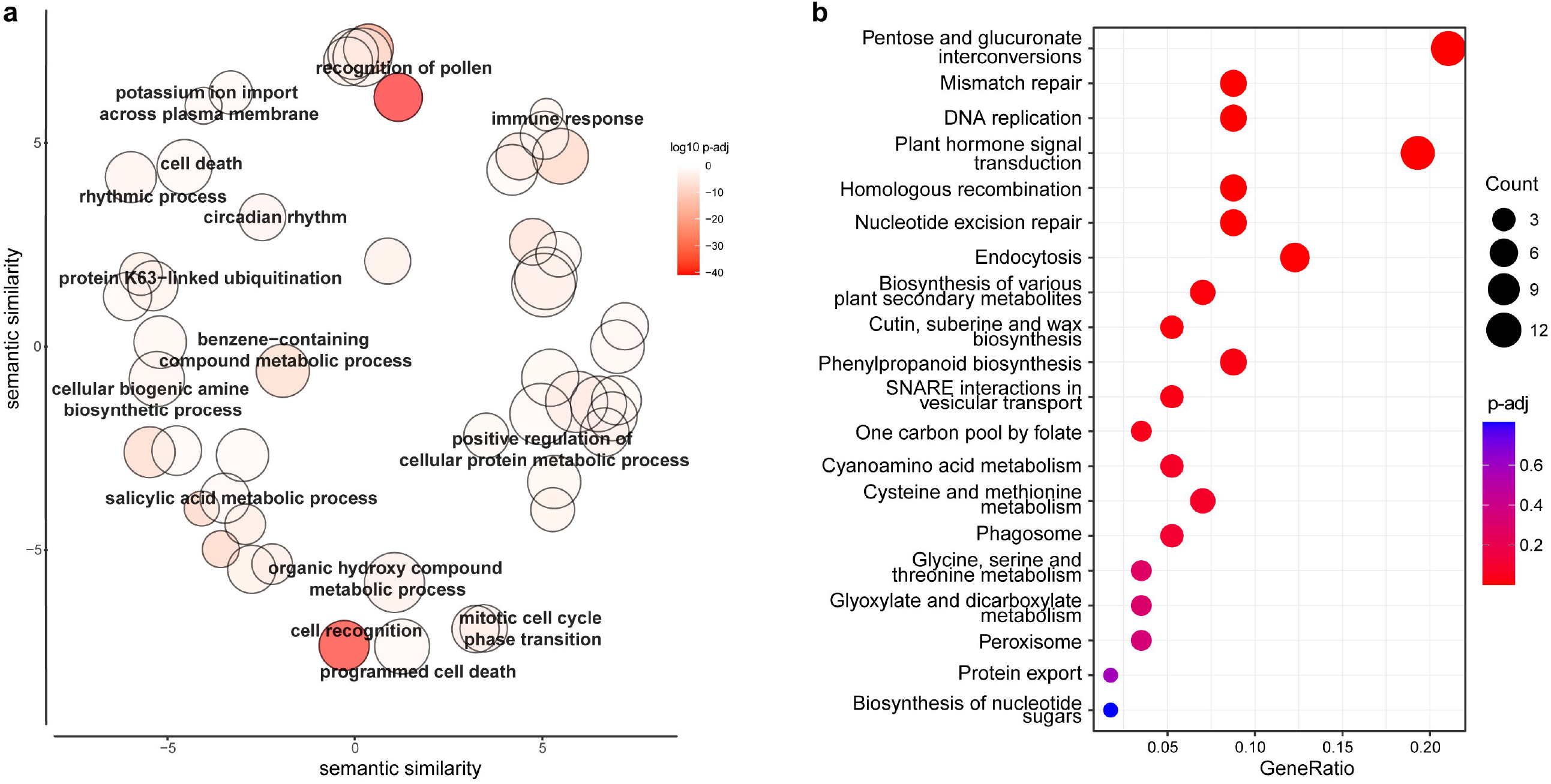
Function enrichment results of gene families undergoing convergently evolutionary rate acceleration in alpine plant genomes. (a) REVIGO clusters of significantly enriched GO terms. Each bubble represents a summarized GO term from the full GO list by reducing functional redundancies, and their closeness on the plot reflects their closeness in the GO graph, i.e., the semantic similarity. (b) Top 20 enriched KEGG pathways. “p-adj” refers to the adjusted *p*-value using the Benjamini– Hochberg method.

#### Self-incompatibility system

Self-incompatibility (SI) in many flowering plants is controlled by the *S* (sterility) locus (Takayama and Isogai 2005). The loss of SI genes in *A. thaliana* is responsible for the evolutionary transition to the self-fertile mating system (Sherman-Broyles et al. 2007). Although self-fertilization is often thought to lead to a decreased fitness of homozygous offspring, this mode ensures reproduction in the absence of pollinators or suitable mates, and therefore can be advantageous for plants to occupy niches in harsh environments (Goodwillie et al. 2005). Our results revealed the biggest orthogroup (OG0000000), which includes the *Arabidopsis* S-receptor kinase (*SRK*) genes and the *S*-locus flanking gene *ARK3* (*RECEPTOR KINASE 3*), undergoing evolutionary rate acceleration and convergent AA shifts in alpine species (fig. 3). GO analysis showed that these genes were functionally enriched in the process of reproduction (recognition of pollen, recognition of pollen, and pollination) and immune system (immune response). Furthermore, we identified the functional domain using NCBI’s conserved domain database (CDD) (Lu et al. 2020). The result showed that these proteins contained the *S*-locus glycoprotein domain (Pfam00954), confirming their functions in SI system.

**FIG. 3.**
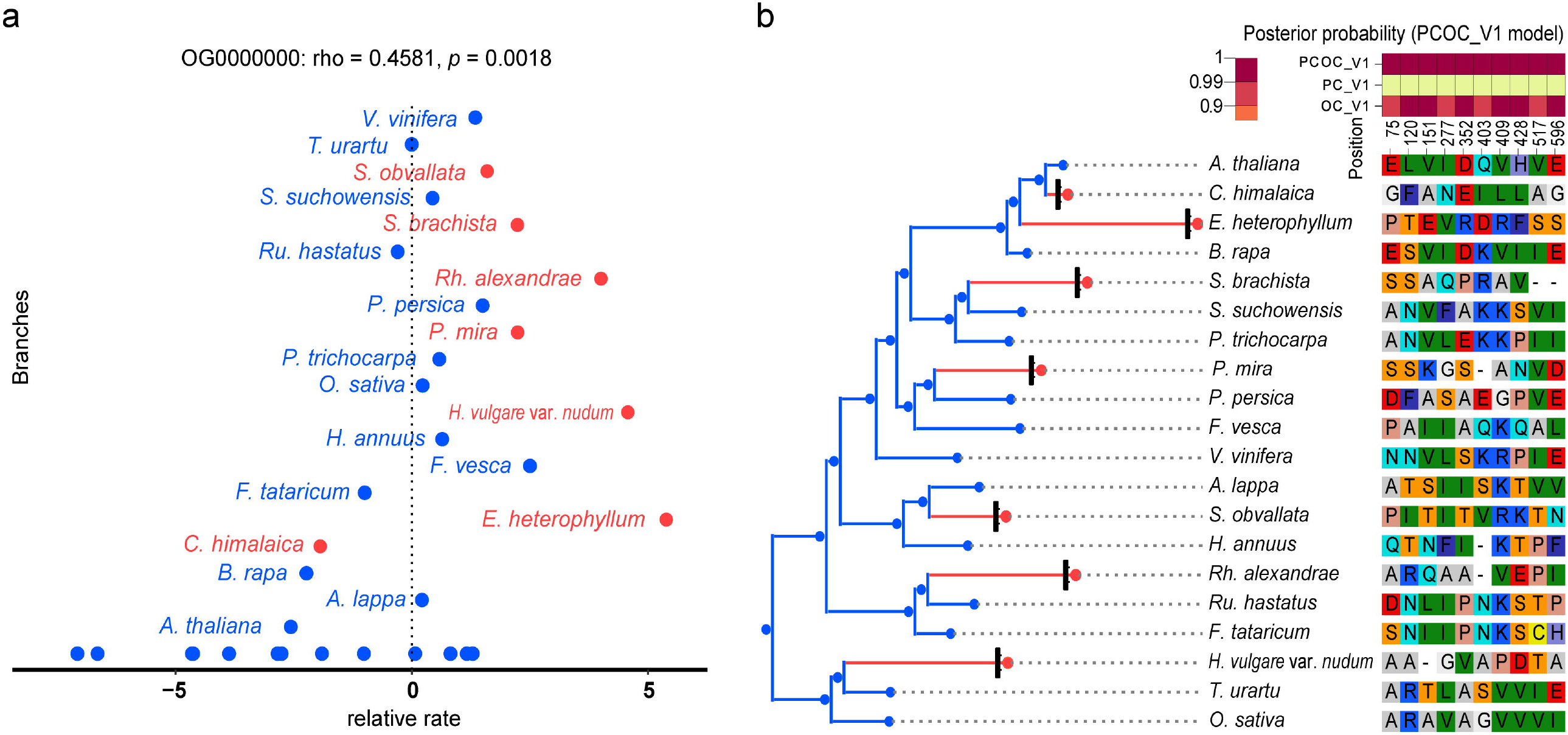
Molecular convergence of *S*-locus genes. (a) Convergent evolutionary rate shifts detected by RERconverge using correlations of relative rates of the gene with traits of interest (here alpine environments) (Kowalczyk et al. 2020). (b) Site-based estimation of convergent amino acid evolution using PCOC method. Only sites with a posterior probability above 0.9 according to the PCOC model are represented. Detailed information for the PCOC, PC, and OC models is described in Rey et al. (2018). Alpine species are in red and their low-land relatives are in blue.

Loss of function at the *S*-locus in alpine *Crucihimalaya* genomes possibly due to relaxed selection were reported (Zhang et al. 2019; Feng et al. 2022), with a similar phenomenon found in the high-altitude Andes maca (*Lepidium meyenii*) genome (Zhang et al. 2016). Our branch-site tests did not detect any signatures of positive selection on these genes, suggesting that acceleration of evolutionary rates may be the result of relaxed constraints. Therefore, we tested for the relaxation of selection on the *S*-locus genes using the RELAX model (Wertheim et al. 2015). RELAX analysis estimated the relaxation intensity parameter *k, k* > 1 indicates intensified selection (i.e., positive or purifying selection) and *k* < 1 suggests relaxed selection. The results showed that five (*C. himalaica, E. heterophyllum, H. vulgare* var. *nudum, S. brachista* and *S. obvallata*) of the seven alpine plants exhibited significantly relaxed selection on *S*-locus genes (*p*-value < 0.05; supplementary table S22). Bingham and Ort (1998) reported that low levels of insect diversity, abundance and activity often occur in alpine ecosystems and hypothesized that these factors may limit the pollination of alpine plants. Species living in isolated habitats like alpine environments or ocean islands are thus less likely to be SI, in line with ‘‘Baker’s Law’’, which assumes that pollen limitation may be an important force driving the transition of mating systems (Cheptou 2012). Pollination biology studies have shown several cases of autonomous selfing in various taxonomically distant species within HHM communities, although the proportion of self-pollinated species has not yet been calculated to test the hypothesis (Sun et al. 2014). We here hypothesize that convergent acceleration of evolutionary rates of *S*-locus genes due to relaxed selection can be evidence of the evolutionary transition from self-incompatibility to the self-compatibility mating system, which is potentially a convergently adaptive process for alpine plants to facilitate their reproduction and the occupation of alpine niches.

#### Cell wall modification

The cuticle membrane lies over and merges into the outer wall of epidermal cells (Martin and Juniper 1970). The primary role of the cuticle, composed of cutin and cuticular waxes, is to mitigate water loss and excessive UV radiation by functioning as a physical barrier between the plant surface and its external environment (Kerstiens 1996). Cuticular waxes are composed of a variety of organic solvent-soluble lipids, consisting of very-long-chain (VLC) fatty acids and their derivatives, as well as secondary metabolites like flavonoids (Pollard et al. 2008). The cell wall modification pathway was found significantly enriched in genes undergoing convergent positive selection (supplementary table S18). Additionally, in the examination of convergent changes in gene family number, we found the most significantly enriched KEGG pathway of co-expanded genes to be cutin, suberine and wax biosynthesis (supplementary table S14), corresponding to the enriched suberin biosynthetic process in the GO analysis (supplementary table S13), including genes encoding fatty acyl-CoA reductases (FARs). FARs catalyze the formation of fatty alcohols, which are common components of plant surface lipids (i.e., cutin, suberin, and associated waxes). We did not detect convergent acceleration of the evolutionary rate of FARs, suggesting possible modifications of FARs through the increase in gene copy number. The results showed that an aldehyde decarbonylase enzyme CER1 underwent a convergent acceleration of evolutionary rate in alpine plants.

Overexpression of the *A. thaliana CER1* gene was reported to promote wax VCL alkane biosynthesis and influences plant response to biotic and abiotic stresses (Bourdenx et al. 2011). The convergent evolution of genes involved in the cutin, suberine and wax biosynthesis implied that cuticular waxes may function as protective screens against UV radiation to protect anatomical structures and dissipate excess light energy for alpine plants.

In addition, we detected positive selection and molecular convergence of three MYB transcription factors, including MYB27, MYB48, and MYB59 (supplementary table S17). Among them, MYB27 was reported to play a role in regulating the accumulation of anthocyanins (Albert et al. 2014), a class of flavonoids. In the STRING database (Szklarczyk et al. 2021), MYB48 was predicted to interact with proteins that are involved in flavonoid biosynthesis, including F3H (naringenin,2-oxoglutarate 3-dioxygenase), catalyzing the 3-beta-hydroxylation of 2S-flavanones to 2R,3R-dihydroflavonols which are intermediates in the biosynthesis of flavonols and anthocyanidins, FLS1 (flavonol synthase/flavanone 3-hydroxylase), catalyzing the formation of flavonols from dihydroflavonols, DFR (dihydroflavonol reductase), catalyzing the conversion of dihydroquercetin to leucocyanidin, and TT5, a member of chalcone-flavanone isomerase family protein (supplementary fig. S15, supplementary table S23). Flavonoids function as antioxidants that reduce DNA damage induced by abiotic stresses such as extreme temperatures, UV radiation and drought, and thus play critical roles in species adapting to high-altitude environments (Agati et al. 2012). Many examples of whole genome studies of alpine plants have shown that expansion and/or positive selection of genes involved in flavonoid biosynthesis constitute an important part of the genomic footprint of alpine adaptation (Zeng et al. 2015; Chen et al. 2019; Wang et al. 2021). Taken together, modifications of the cell wall in alpine plants through evolutionary expansion and adaptive convergence of genes involved in the biosynthesis of cuticular waxes and flavonoids might be vital strategies for adaptation to dramatic weather changes and extensive UV radiation in high altitude.

With the newly generated transcriptomic data of *S. obvallata*, we were able to investigate expression patterns of genes related to the biosynthesis of cuticular waxes and flavonoids. Five tissues with three biological replicates, including three from leaves (basal leaves JL, middle leaves ML, and bract leaves BL) and two from flowers and stems, were sampled (supplementary fig. S16, supplementary table S3). After mapping the RNA-seq data to the assembled genome of *S. obvallata*, 25,096 genes had expression profiles and were retained for differential gene expression (DEG) analysis. The results showed that, compared to basal and middle leaves, bract leaves exhibit 1,071 significant up-regulated genes (fig. 4a). These genes were significantly enriched in cytochrome P450, cutin, suberine and wax biosynthesis and isoflavonoid biosynthesis KEGG pathways (fig. 4b). Furthermore, we analyzed expression profiles of genes involved in the biosynthesis of cuticular waxes and flavonoids. The results showed that many genes had higher expression levels in bract leaves than in other leaf tissues. For example, *CER1, CER3, CER4*, and *MAH1* in the cuticular wax biosynthetic pathway (fig. 4c), and *4CL, CHS, CHI, F3’H, TT7* and *OMT* in the flavonoid biosynthetic pathway (fig. 4d). These results suggest that the accumulation of cuticular waxes and flavonoids is an important genetic pathway from normal leaves to leafy bracts. Our findings provide new insights into the genetic basis of the specialized ‘glasshouse’ morphology for a better understanding of plant morphological adaptation.

**FIG. 4.**
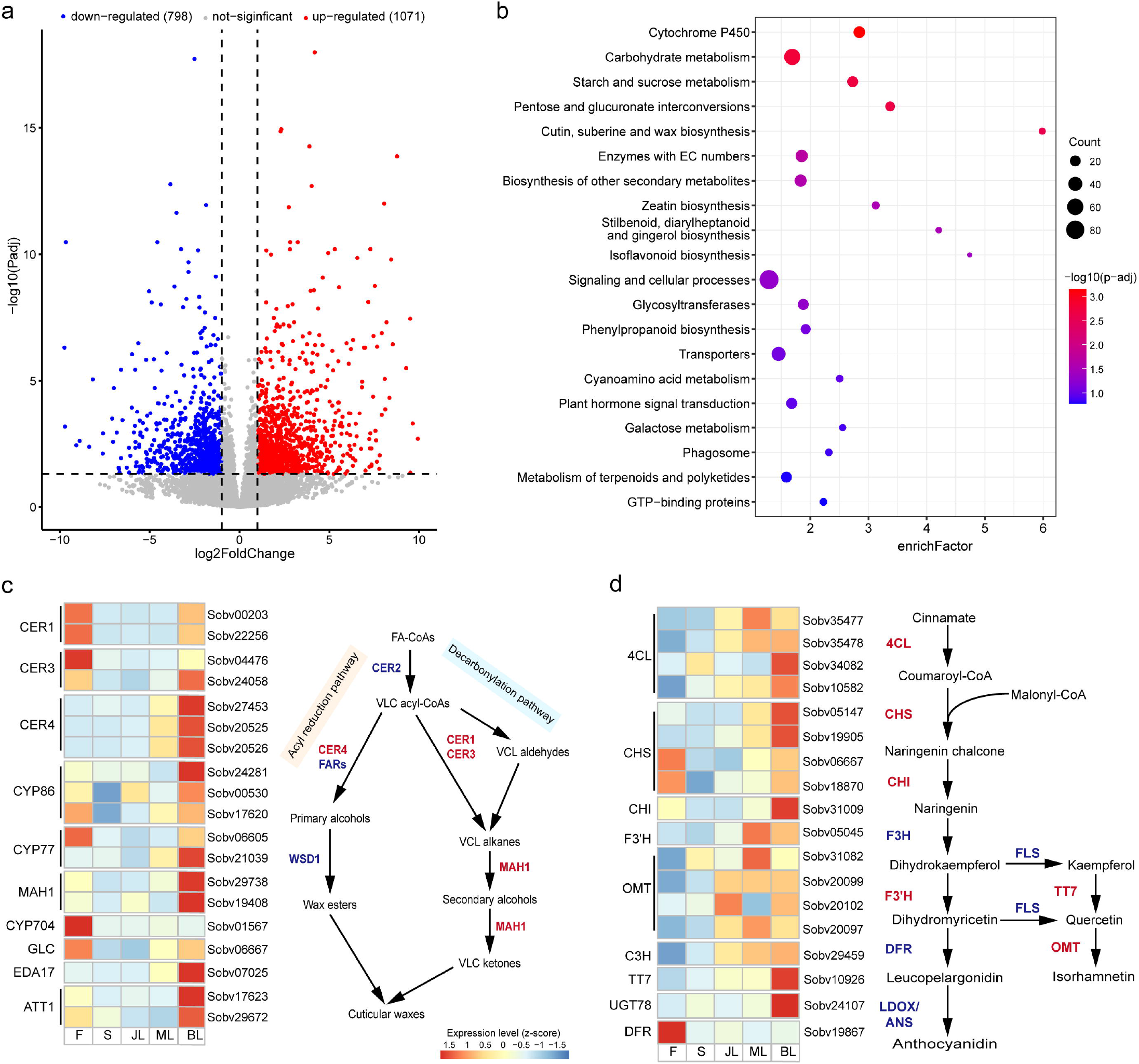
Highly expressed genes in bract leaves of *Saussurea obvallata*. (a) Differentially expressed genes between bract leaves with normal leaves. (b) Top 20 KEGG pathways of significantly up-regulated genes in bract leaves. “p-adj” refers to the adjusted *p*-value using the Benjamini–Hochberg method. Expression profiles of genes involved in cuticular wax (c) and flavonoid biosynthetic pathways (d) in different tissues of *S. obvallata* (F: flowers, S: stems, JL: basal leaves, ML: middle leaves and BL: bract leaves). High expressed genes in BL are shown in red in the simplified pathway models. The bar represents the gene expression level of each gene (z-score). Abbreviations: FA-CoA, fatty acyl-coenzyme A; VLC, very-long-chain; CER, protein eceriferum; FAR, fatty acid synthetase; WSD1, diacylglycerol acyltransferase 1; MAH1, midchain alkane hydroxylase 1; 4CL, 4-coumarate: CoA ligase; CHS, chalcone synthase; CHI, chalcone isomerase; F3H, flavanone 3-hydroxylase; F3’H, flavonoid 3’-hydroxylase; FLS, flavonol synthase; TT7, transparent testa 7; OMT, O-methyltransferase; DFR, dihydroflavonol 4-reductase; LDOX/ANS, leucoanthocyanidin dioxygenase/anthocyanidin synthase. The pathways for the cuticular wax biosynthesis was adapted from the study of Wang et al. (2020), and for flavonoid biosynthesis was adapted from the study of Yonekura-Sakakibara et al. (2008).

#### DNA repair and stress resistance pathways

The hypoxia and intense UV radiation in alpine environments exert highly abiotic stress that can cause DNA, RNA, and protein damage. DNA repair processes thus play an important role in the high-altitude adaptation of plants, similar to evidence in alpine animals (Li et al. 2018). The *NBS1* gene, involved in DNA repair, cellular response to DNA damage stimulus and double-strand break repair, was found to be under convergent selection in alpine genomes (supplementary table S17). Moreover, several significantly enriched GO terms related to DNA repair and protein ubiquitination were detected undergoing convergent evolution in alpine plants (fig. 2a, supplementary table S17). Similar KEGG pathways were also identified, including mismatch repair, homologous recombination and nucleotide excision repair (fig. 2b, supplementary table S18). Examples included the *UEV1* genes, enriched in protein K63-linked ubiquitination pathway that reportedly play a role in DNA damage responses and error-free post-replicative DNA repair by participating in lysine-63-based polyubiquitination reactions (Wen et al. 2008), and genes encoding OB-fold proteins which were found to be important for genomic stability including DNA replication, recombination, repair, and telomere homeostasis (Flynn and Zou 2010).

In addition to low oxygen levels and excessive UV radiation, high-altitude environments pose various threats to living organisms from an unpredictable climate. We found many genes with convergent evolutionary rate shifts were significantly enriched in stress resistance pathways, such as genes involved in the hormone signal transduction (cell recognition, intracellular signal transduction, and cytokinin-activated signaling pathway), the cell rhythm system (circadian rhythm, cell cycle phase transition, and programmed cell death), and the regulation of enzyme activity pathways (regulation of kinase activity, regulation of GTPase activity, and regulation of transferase activity) (fig. 2a, supplementary tables S17). Also, some KEGG pathways were found to be significantly enriched in metabolic pathways that may contribute to stress resistance, such as plant hormone signal transduction, phenylpropanoid biosynthesis, and glycine, serine, and threonine metabolism (fig. 2b, supplementary table S18). Plant hormones like cytokinin and ethylene play pivotal regulatory roles in plant growth and development, including cell division, shoot initiation, light responses, and leaf senescence. We found that Type-A *Arabidopsis* response regulator (ARR) protein family, involved in response to cytokinin and cytokinin signal transduction, undergoes molecular convergence in alpine plants. An experimental study revealed that *arr* mutants show altered red-light sensitivity, indicating an important role of type-A ARRs in light adaptation (To et al. 2004). The circadian clock was selected as a mechanism in the control of cell-cycle progression to avoid sunlight-induced DNA damage in ancient unicellular organisms (Hut and Beersma 2011). Circadian rhythm in plants regulates multiple processes such as photosynthesis, flowering, seed germination, and senescence (Srivastava et al. 2019). Genes involved in the regulation of the rhythm system were found to be convergently accelerated evolutionary rates in alpine plants, such as the transcription factor *MYB59*, which participates in the regulation of the cell cycle, mitosis and root growth by controlling the duration of metaphase. The *myb59* mutant was found to have longer roots, smaller leaves and smaller cells than wild-type plants (Fasani et al. 2019), which are commonly observed in alpine plants. Taken together, these results suggested that adaptation to high altitude requires the participation of multiple biological processes.

## Conclusions

Taking advantage of an integrative genomic data set of alpine plants, our study unraveled the genomic underpinnings of convergent adaptation to high-altitude environments (fig. 5). We identified convergently expanded gene families involved in the oxygen-level response and contracted gene families related to disease resistance in alpine plants, potentially reflecting adaptations to high-altitude environments experiencing hypoxia and depauperate pathogen communities. Our results showed that alpine plants have undergone convergent molecular evolution of genes involved in reproduction and development, self-incompatibility, cell wall modification, as well as DNA repair and stress resistance pathways that may contribute to the survival in extremely cold, high UV radiation and hypoxia environments. Both molecular convergence in changes of gene copy number and accelerated evolutionary rates are consequences of selective pressures posed by the surrounding environment. Thus, genomic signatures of convergent evolution detected here are direct evidence for alpine plants associated with independently colonizing, evolving and adapting to cold, high UV radiation and hypoxia environments of the HHMs. The alpine plants included in this study belong to taxonomically distant plant orders, hence, standing genetic variation and localized introgression of regions of the genome can be ruled out as probable causes of genomic convergence. Identical *de novo* mutations must therefore have occurred independently in each taxon during evolution from a low-altitude ancestor. Furthermore, we gain valuable insights into genetic bases underlying the adaptive ‘greenhouse’ morphology by mining differentially expressed genes involved in the biosynthesis of cuticular waxes and the accumulation of flavonoids that might be vital strategies for adaptation to dramatic weather changes and extensive UV radiation in high altitudes. Taken together, our study reveals genomic convergence underlying adaptation to alpine environments at a high-taxonomic level of angiosperms, providing a framework for further understanding of high-altitude adaptation. Further genomic study of alpine adaptation should provide detailed evidence related to morphological and physiological specializations using updated data, such as proteome and metabolome, while also referring to discoveries from phylogenetically distant taxon to evaluate the evolutionary convergence in similar environments.

**FIG. 5.**
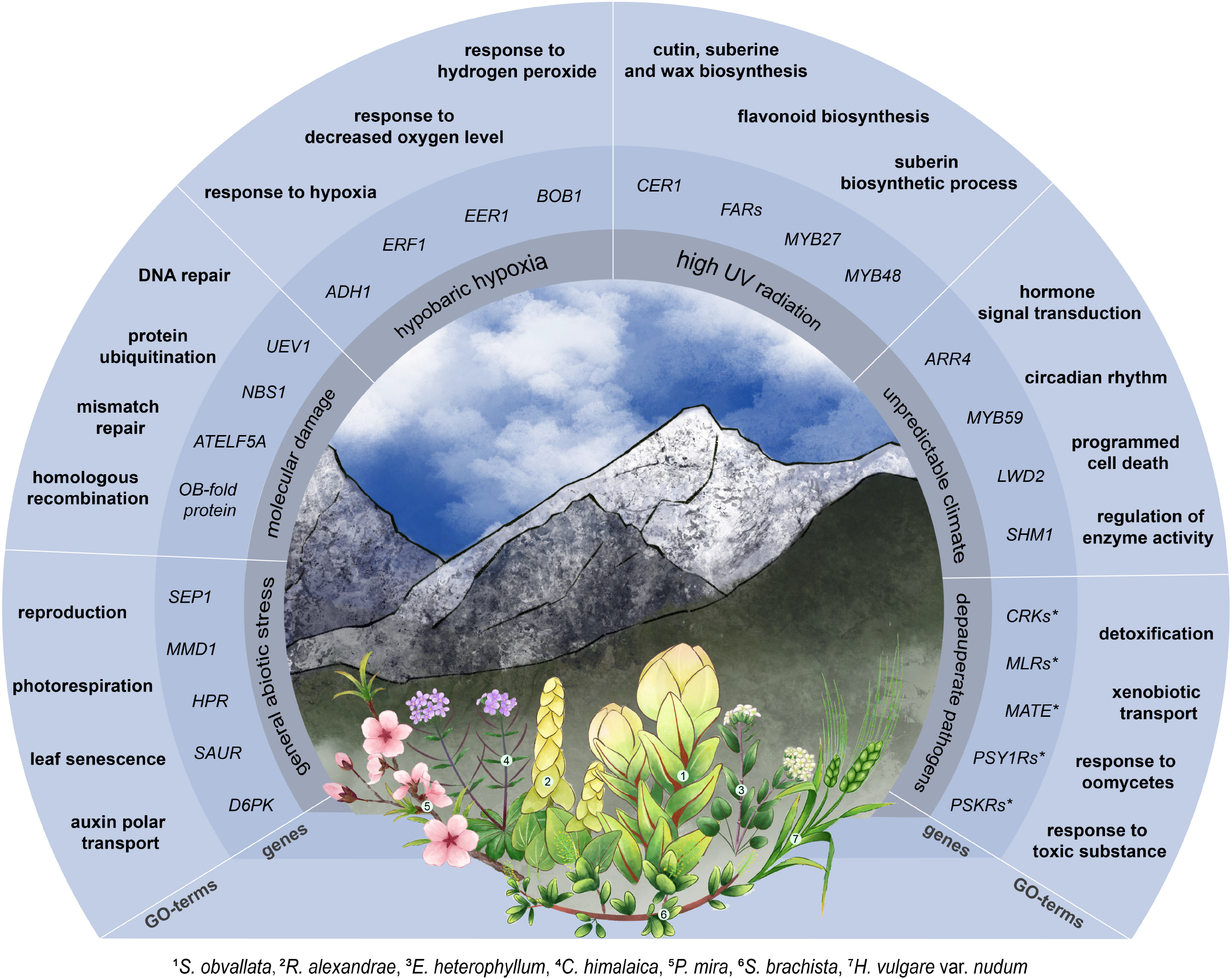
Summary of convergent adaptation to high-altitude environments for seven alpine species. The outer circle shows examples of enriched GO-terms (biological pathways). The middle circle shows examples of candidate genes named after the *A. thaliana* orthologs. The inner gray circle shows environmental stresses from high altitudes. Species names are provided below the picture. *Genes undergone convergent contraction.

## Materials and Methods

### Plant material, DNA extraction and sequencing

Fresh leaves of wild *S. obvallata* and *R. alexandrae* individuals were collected from Songpan county, Sichuan Province, China (102°45′E,30°23′N) and Linzhi county, Tibet Province, China (102°45′E,30°23′N), respectively. All samples were sent to Wuhan Benagen Technology Company Limited (Wuhan, China) for genome sequencing. Total genomic DNA was extracted using the Qiagen DNeasy Plant Mini Kit. For the Illumina short reads, DNA libraries with 500 bp insert sizes were constructed and sequenced using an Illumina HiSeq 4000 platform. For Oxford Nanopore sequencing, libraries were prepared using the SQKLSK109 ligation kit using the standard protocol, and the purified library was loaded onto primed R9.4 Spot-On Flow Cells and sequenced using a PromethION sequencer (Oxford Nanopore Technologies, Oxford, UK). The Hi-C sequencing was performed as follows: extracted DNA was first crosslinked by 40ml of 2% formaldehyde solution to capture interacting DNA segments, the chromatin DNA was then digested with the *DpnII* restriction enzyme, and libraries were constructed and sequenced using Illumina HiSeq 4000 instrument with 2×150 bp reads.

For RNA sequencing, fresh tissue samples were collected and immediately frozen in liquid nitrogen. Three biological replicates of five tissues of *S. obvallata* were sampled. Total RNA was extracted using the TRIzol® Reagent (Invitrogen, Shanghai, China). Paired-end cDNA libraries were constructed using TruSeq Stranded mRNA Library Prep Kit (Illumina), and were sequenced using Illumina HiSeq 4000 platform.

### Genome assembly and quality control

Before genome assembly, Kmerfreq (https://github.com/fanagislab/kmerfreq) was used for counting *K*-mer frequency with the *K*-mer set to 19, and GCE v1.0 was used for estimating the genome size (Liu et al. 2013). The ONT long reads were corrected and assembled using NextDenovo v2.3.1 (https://github.com/Nextomics/NextDenovo) with default parameters. The contigs were polished using NextPolish v1.4.1 (Hu et al. 2020) with the long reads and Pilon v1.23 (Walker et al. 2014) with Illumina short reads for three rounds. Hi-C data were used to execute chromosome conformation using the 3D-DNA pipeline with default parameters. The accuracy of Hi-C based chromosomal assembly was improved using Juicerbox’s chromatin contact matrix (Dudchenko et al. 2017). The completeness and continuity of the assemblies were evaluated by the statistics of BUSCO and LAI, respectively.

### Genome annotation

#### Repeat annotation

TEs were identified based on *de novo* and homology-based strategies. RepeatMasker v4.0.7 was used to run a homology search for known repeat sequences against the Repbase database v22.11 (Tarailo-Graovac and Chen 2009). RepeatModeler v2.0.10 was employed to predict the TEs based on the *de novo* method (Flynn et al. 2020). LTRharvest v1.5.10, LTR_FINDER v1.05 and LTR_retriever v1.8.0 were used to build an LTR library with default parameters (Xu and Wang 2007; Ellinghaus et al. 2008; Ou and Jiang 2018). Finally, RepeatMasker was used to merge these library files of the two methods and to identify the repeat contents.

#### Gene prediction

A combination of *de novo*-, homology-, and transcript-based methods were used for gene prediction in both genomes. RNA-seq reads were assembled using Trinity v2.1.1 using the *de novo*-based and genome-guided modes, respectively. Coding DNA sequences (CDS) and protein sequences were predicted with TransDecoder (http://transdecoder.github.io). Homologs were predicted by mapping protein sequences using GeMoMa v1.6.1 (Keilwagen et al. 2016). Sequences of *Arabidopsis thaliana, Oryza sativa* and *Solanum tuberosum* were mapped to both genomes. Additionally, sequences of *Helianthus annuus, Lactuca sativa, Cynara cardunculus* and *Mikania micrantha* were mapped to *S. obvallata*, and sequences of *Fagopyrum tataricum* and *Rumex hastatus* were mapped to *R. alexandrae*, respectively. A *de novo* gene prediction was performed with Braker2 v2.1.5 and GlimmerHMM v3.0.4 (Majoros et al. 2004; Bruna et al. 2021). Assembled transcripts were used for training gene models in Braker2. Gene models from the three main sources were merged to produce consensus models using EVidenceModeler v1.1.1 (Haas et al. 2008).

#### Gene functional annotation

The annotated protein-coding genes were used for a BLAST search against the UniProt and NCBI nonredundant protein databases to predict gene functions. The functional domains of protein sequences were identified by InterProScan v5.51-85.0 using data from Pfam (Jones et al. 2014). KEGG and gene ontology (GO) terms of the gene models were obtained using eggNOG-mapper v2.0.1 (Cantalapiedra et al. 2021).

### Orthogroup inference and alignment

Protein sequences from the seven alpine genomes as well as 13 relatives living at low altitudes were used for subsequent comparative analyses. OrthoFinder v2.5.4 was used to construct orthogroups for all species with default settings (Emms and Kelly 2019). Because the included species are phylogenetically distant (widely dispersed across the eudicots and monocots), OrthoFinder recovered an extremely low number of strictly single-copy orthogroups. We therefore reduced orthogroups that contained multiple gene copies per species to one copy per species based on the smallest genetic distance following the study of Birkeland et al. (2020). Briefly, all orthogroups were aligned based on protein sequence using MAFFT v7.3 (Katoh and Standley 2013), and genetic distances between all pairs of genes were calculated as Kimura protein distances (Kimura 1980). One gene copy per species was retained based on the smallest genetic distance to the longest protein sequence of *A. thaliana* in each orthogroup. The rationale is that the annotation of *A. thaliana* genome is complete and reliable, and our subsequent functional analyses of candidate genes were based on the *A. thaliana* orthologs. The resulting orthogroup sequences that contained only one protein sequence per species were realigned using PRANK v170427 (Löytynoja 2014). Coding sequences of genes in orthogroups were extracted based on the same gene identifier of protein sequences and were aligned using PRANK with the codon mode.

### Phylogeny and estimation of divergence time

Protein sequences of 195 single-copy orthogroups were used for phylogenetic inference with RAxML v8.2.12 using the PROTGAMMAAUTO substitution model with 500 bootstrap replicates (Stamatakis 2014). Divergence time was estimated in the MCMCtree program from PAML v4.9j (Yang 2007), using the approximate likelihood method with the tree topology inferred with RAxML, an independent substitution rate (clock = 2), and the HKY85+GAMMA model. Three calibration points were assigned based on the TimeTree database (Kumar et al. 2017) (http://www.timetree.org/, accessed on 1 May 2022): the MRCA (most recent common ancestor) of rosids (95% HPD 105 - 115 Mya), the MRCA of *Pentapetalae* (95% HPD 110 - 124 Mya), and the MRCA of *Mesangiospermae* (95% HPD 148 - 173 Mya). Samples were drawn every 1,000 MCMC steps from a total of 10^6^ steps, with a burn-in of 10^5^ steps. Convergence was assessed by comparing parameter estimates from two independent runs, with all effective sample sizes >200.

### Gene family evolution

The change in gene family number was examined using CAFÉ5 (Mendes et al. 2020). Besides the base model, the number of gamma categories (-*k*) was set to estimate separate lambda values for different lineages in the tree (Gamma model). The highest likelihood was found using *k* = 2 rate categories (-lnL = 286114), with λ = 0.00553 and α = 1.58. Gene family expansions or contractions were identified only when the change in gene count was significant with a *p*-value < 0.05. Genome-wide NBS-LRR genes were manually identified using HMMER v3.2 with an e-value 1e-05 (Wheeler and Eddy 2013). The NBS-LRR protein domains (NB-ARC: PF00931; RPW8: PF05659; TIR: PF01582; LRR: PF00560, PF07723, PF07725 and PF12799) were retrieved from Pfam (http://pfam.xfam.org) and were used to identify conserved motif of NBS-LRR genes in sampled genomes. The ML phylogenetic tree of NBS-LRR was constructed using RAxML. Then, Notung v2.9 was used to recover gains and losses of NBS-LRR genes by reconciling the NBS-LRR gene tree (Chen et al. 2000). The concatenated tree reconstructed by RAxML was used as input topology.

### Functional enrichment analysis

GO and KEGG over-representation tests were performed using clusterProfiler v4.3.4 implemented in R to identify significantly enriched pathways (Wu et al. 2021). GO and KEGG terms were assigned according to the orthologous genes of *A. thaliana* genome. In the ‘enrichGO’ function, we set ‘ont = BP’ to only search for enriched biological pathways (BP). The resulting *p-*values were corrected for multiple comparisons using a Benjamini–Hochberg FDR correction. A criterion of *p*-adjust < 0.05 was used to access the significance of enrichment analyses.

### Tests for positive selection and relaxation of selection

The branch-site model implemented in CodeML (PAML package) was performed to test for positive selection. In this test, an alternative model allowing sites to be under positive selection on the foreground branch was contrasted to a null model limiting sites to evolve neutrally or under purifying selection using a likelihood-ratio test (LRT). LRT *p*-values were computed based on Chi-squared distribution (df = 2) and were corrected for multiple tests at a *p*-adjust < 0.05 threshold using a Benjamini-Hochberg FDR correction. A gene showing signature of positive selection in more than three alpine species was identified as a convergent selected gene. Additionally, RELAX was employed to test for the relaxation of selection on *S*-locus genes using the likelihood ratio test by comparing the model fixing *k* = 1 and the model allowing *k* to be estimated (Wertheim et al. 2015). In both analyses, seven tests were conducted separately by setting each alpine species as the foreground branch.

### Convergent evolutionary rate shifts

To perform the RERconverge analysis, we first used PAML to estimate maximum-likelihood gene trees whose branch lengths represent evolutionary rates using the number of amino acid substitutions. RERs were calculated using ‘getAllResiduals’ function with weight = T, scale = T and cutoff = 0.001 (Kowalczyk et al. 2019). We set alpine species as foreground branches (branches of the tree with the trait of interest) and ran ‘foreground2Paths’ function to estimate RERs for all genes and to correlate them with trait evolution (i.e., alpine habitats in our study). We then used ‘correlateWithBinaryPhenotype’ function to test for a significant association between RERs and traits across all branches of the tree with a *p*-value < 0.05.

### Convergent AA evolution

Two models were compared in PCOC analysis: the convergent model in which a site on convergent branches evolves under a profile different from that of the nonconvergent branches, and nonconvergent (null) model in which a site evolves under a single amino acid profile throughout the phylogeny (Rey et al. 2018). PCOC then detected convergent sites by identifying the better fit between the two models. To filter for only sites with strong evidence for convergent profile shifts, we set a posterior probability threshold of > 0.9 in the analysis.

### Differential gene expression (DEG) analysis

RNA-seq data of *S. obvallata* were mapped to the assembled genome using HISAT2 v2.2.1 (Kim et al. 2019). Only uniquely mapped paired-end reads were retained for read counting by featureCounts v2.0.3 (Liao et al. 2014) to generate the count and Transcripts per Kilobase Million (TPM) tables. DEG analyses among the five tissues were performed in DESeq2 v1.36.0 (Love et al. 2014), with a *p*-value <0.05 as a cut-off and a log2 fold change cut-off of 1.

## Data Availability

The genomic data generated and analyzed in this study including the raw sequencing data of Oxford Nanopore, Illumina, Hi-C and RNA-seq, as well as genome assembly have been deposited in China National GeneBank (CNGB, https://db.cngb.org/) under accession number CNP0003451. All the custom scripts and specific command lines have been deposited at GitHub (https://github.com/ZhangXu-CAS/Alpine_genome).

## Supplementary Material

Supplementary data are available at *Molecular Biology and Evolution* online.

## Acknowledgements

The authors thank Ticao Zhang and Yingxiong Qiu for insightful comments on our paper, and thank Xinyi Guo for providing genome data. This work was supported by the Second Tibetan Plateau Scientific Expedition and Research program (2019QZKK0502), the Strategic Priority Research Program of Chinese Academy of Sciences (XDA20050203), the Key Projects of the Joint Fund of the National Natural Science Foundation of China (U1802232), the Youth Innovation Promotion Association of Chinese Academy of Sciences (2019382), the Yunnan Young & Elite Talents Project (YNWR-QNBJ-2019-033) and the Ten Thousand Talents Program of Yunnan Province (202005AB160005).

## Author contributions

T.D., H.W. and H.S. conceived and led the project. X.Z., T.K., W.D. and Z.Q. processed data, performed analyses and drew figures. X.Z., H.Z., T.F., L.L. and T.D. collected materials. X.Z. wrote the manuscript. J.B.L., Y.S., J.H., T.D., H.W. and H.S. revised the manuscript.

