## Supplementary files for "Genomic underpinnings of convergent adaptation to high altitudes for alpine plants"

*Zhang et al.*

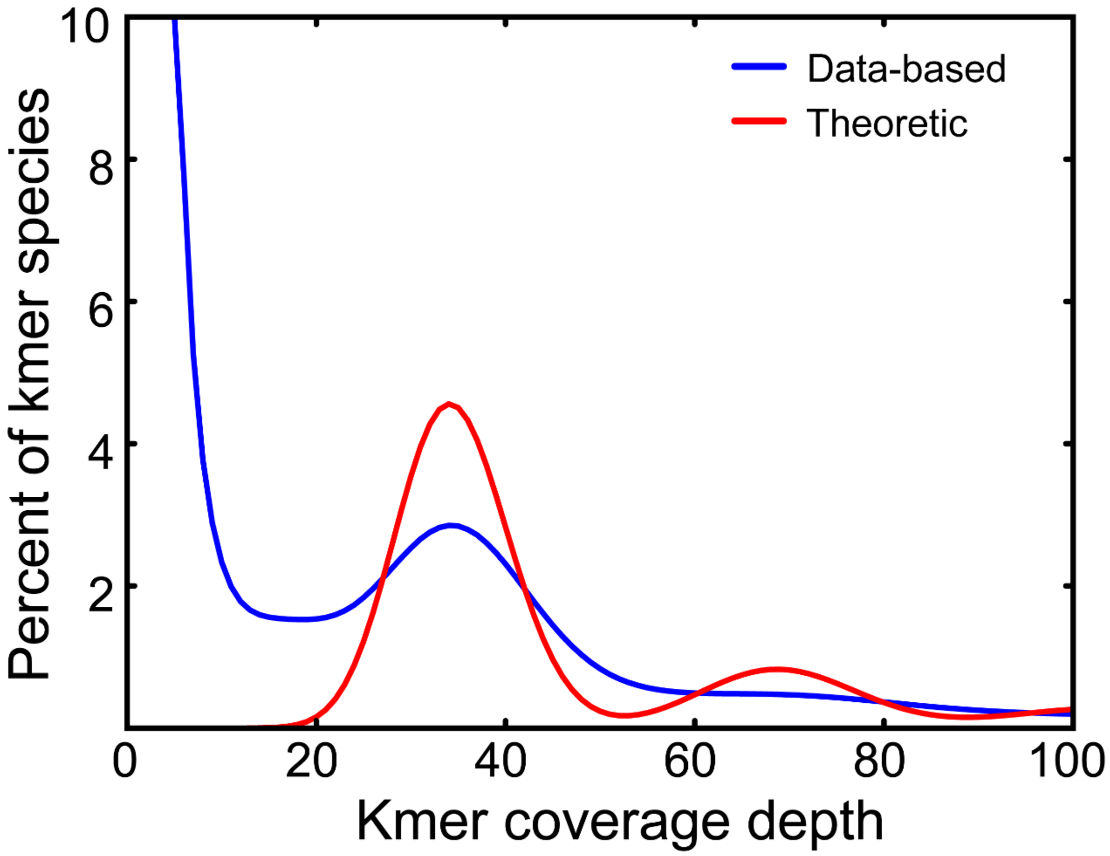


**Fig. S1. *K*-mer based analysis to estimate the genome size of *S. obvallata*.**

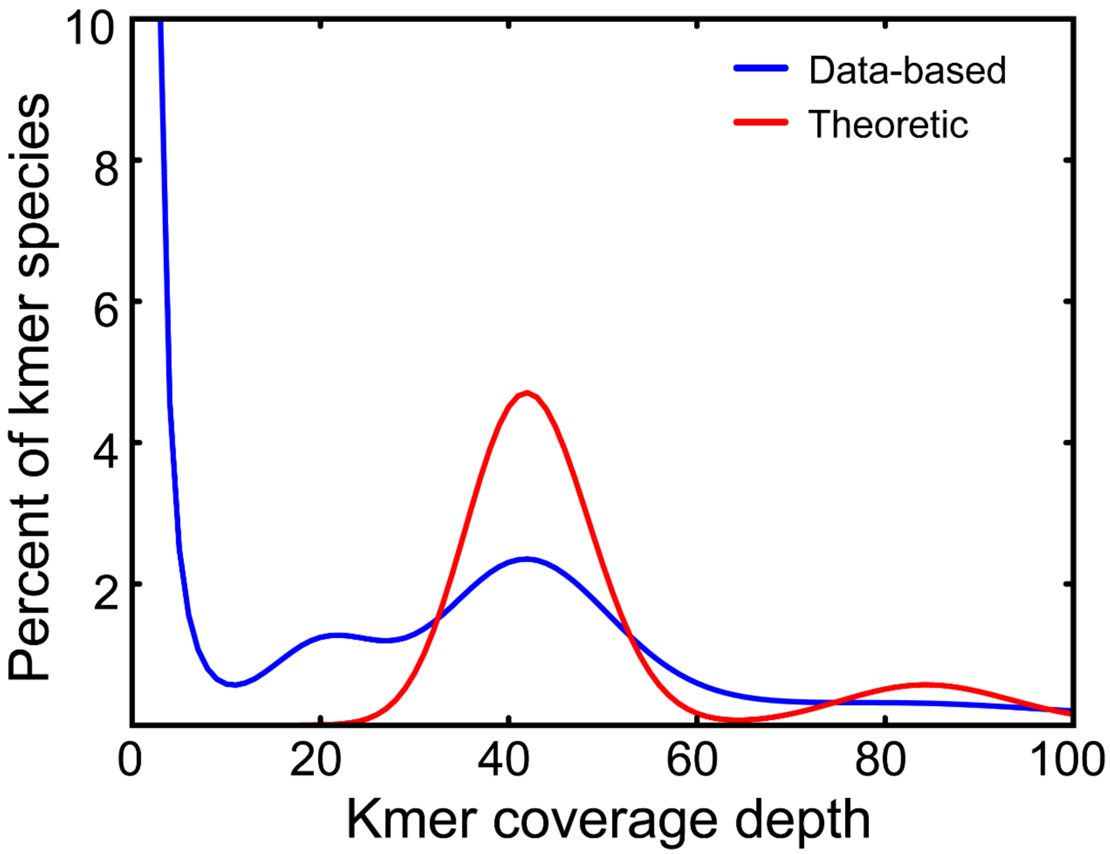


**Fig. S2. *K*-mer based analysis to estimate the genome size of *R. alexandrae*.**

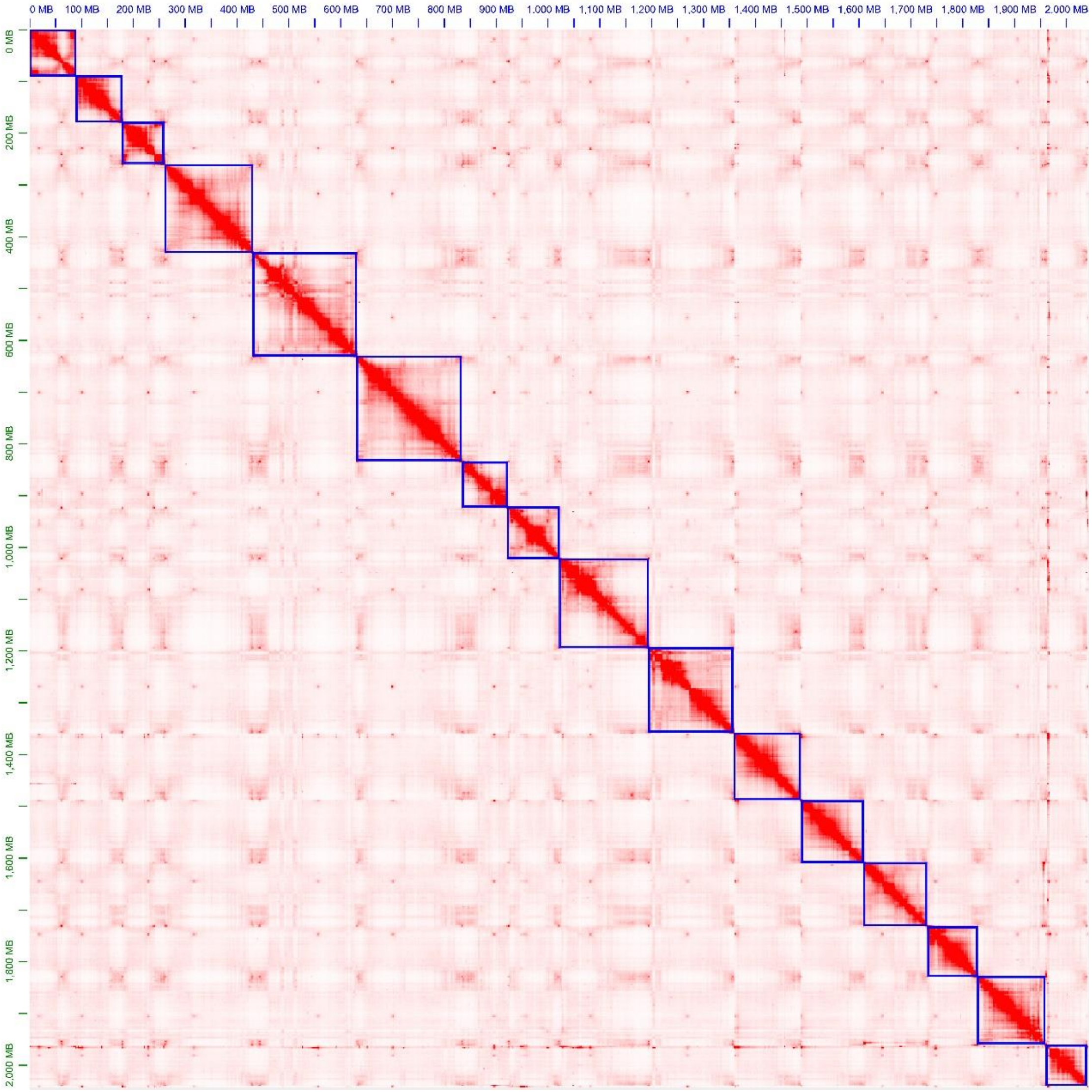


**Fig. S3. The Hi-C assisted assembly of *S. obvallata* pseudomolecules.** Heatmap showing Hi-C interactions with darker red color indicating a higher contact probability.

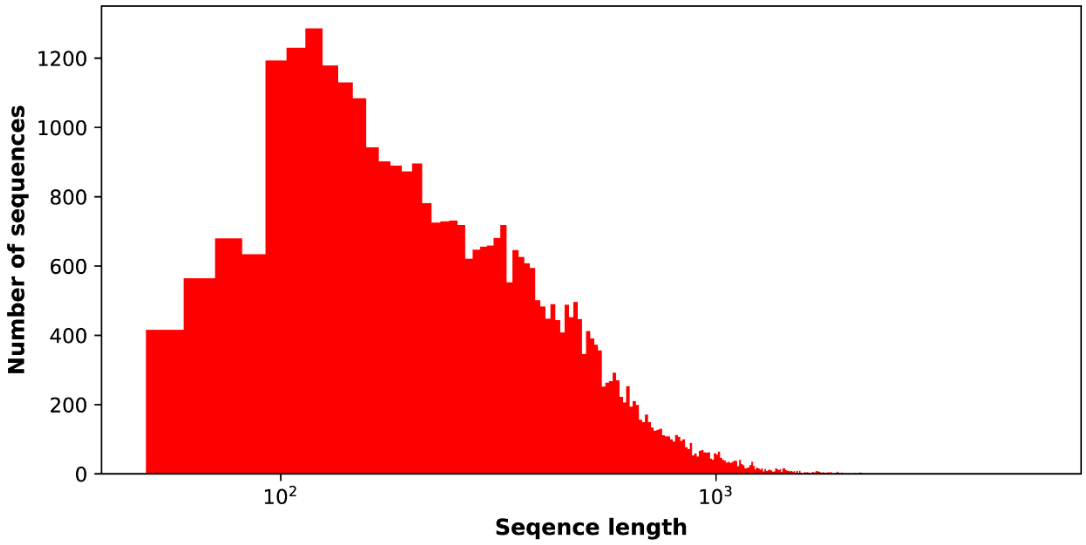


**Fig. S4. The distribution of sequence length of predicted protein-coding genes in *S. obvallata* genome.**

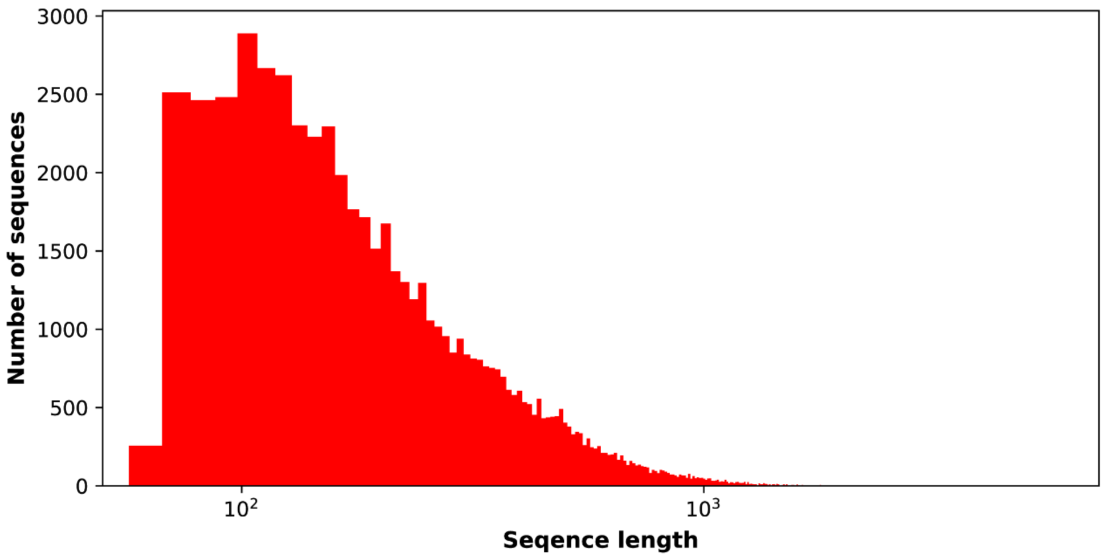


**Fig. S5. The distribution of sequence length of predicted protein-coding genes in *R. alexandrae* genome.**

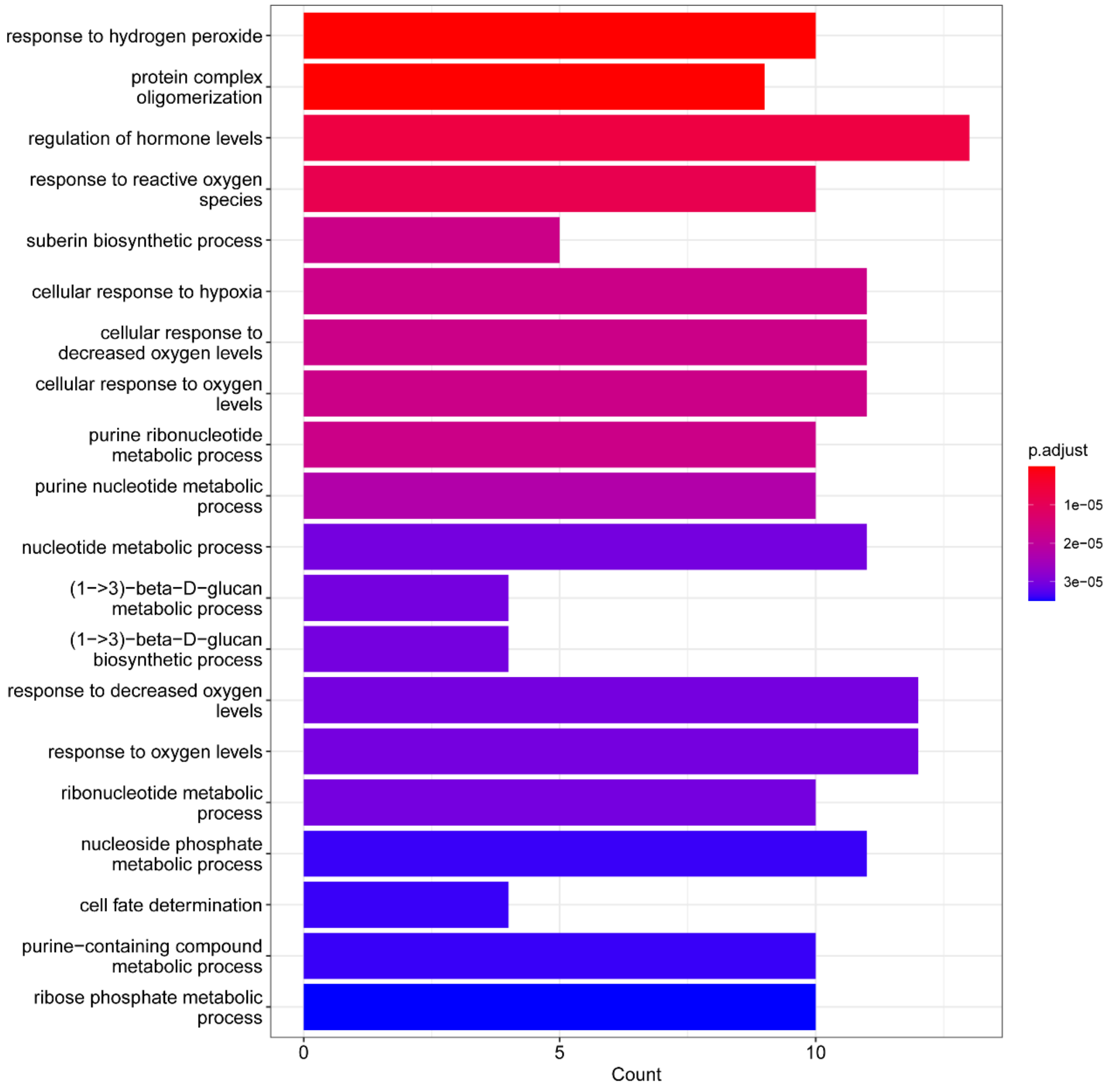


**Fig. S6. Top 20 enriched GO terms of convergently expanded gene families in alpine plant genomes.** “p. adjust” refers to the adjusted *p*-value of the Benjamini–Hochberg false discovery rate (FDR), and the same below.

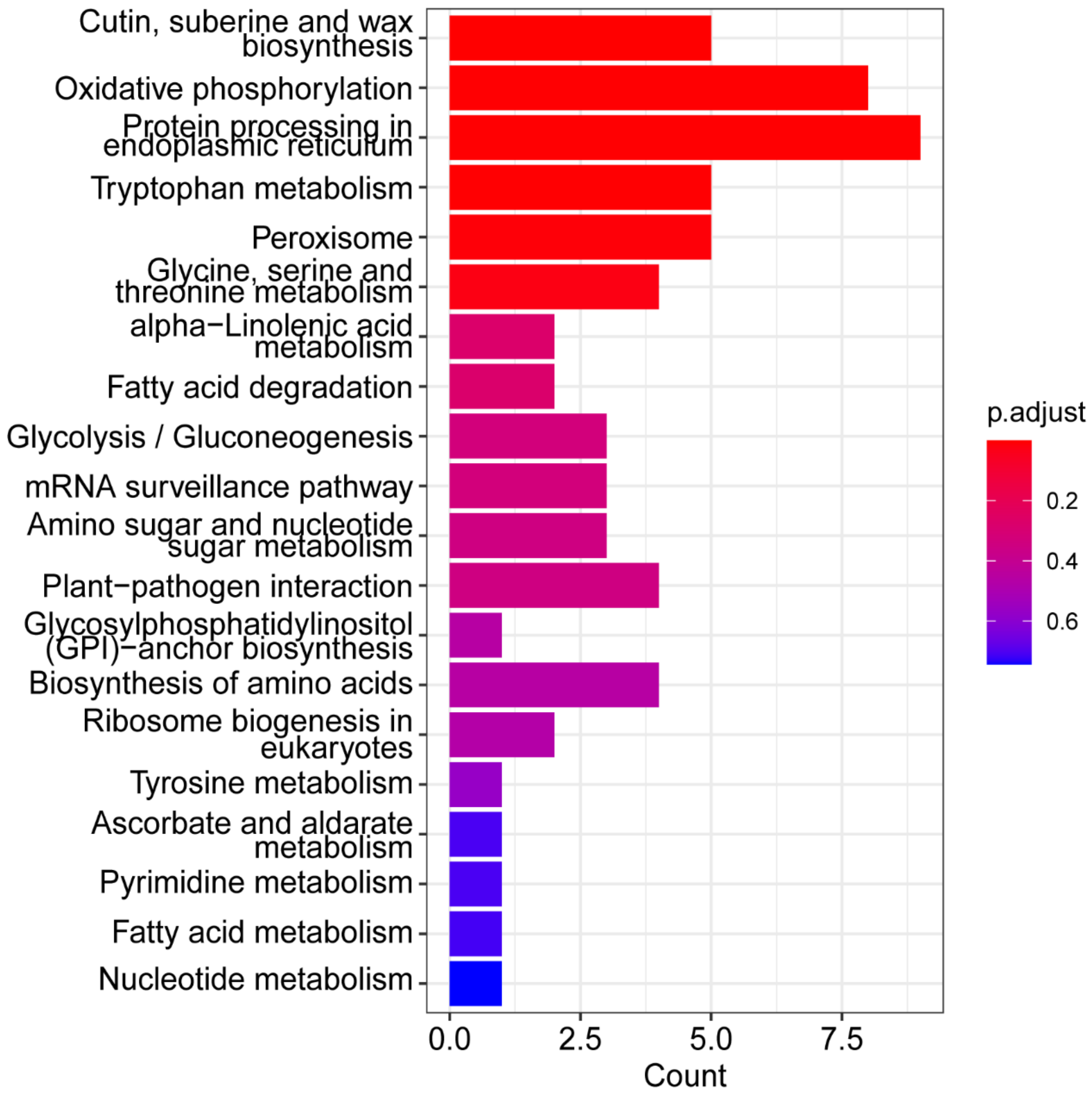


**Fig. S7. Top 20 enriched KEGG pathways of convergently expanded gene families in alpine plant genomes.**

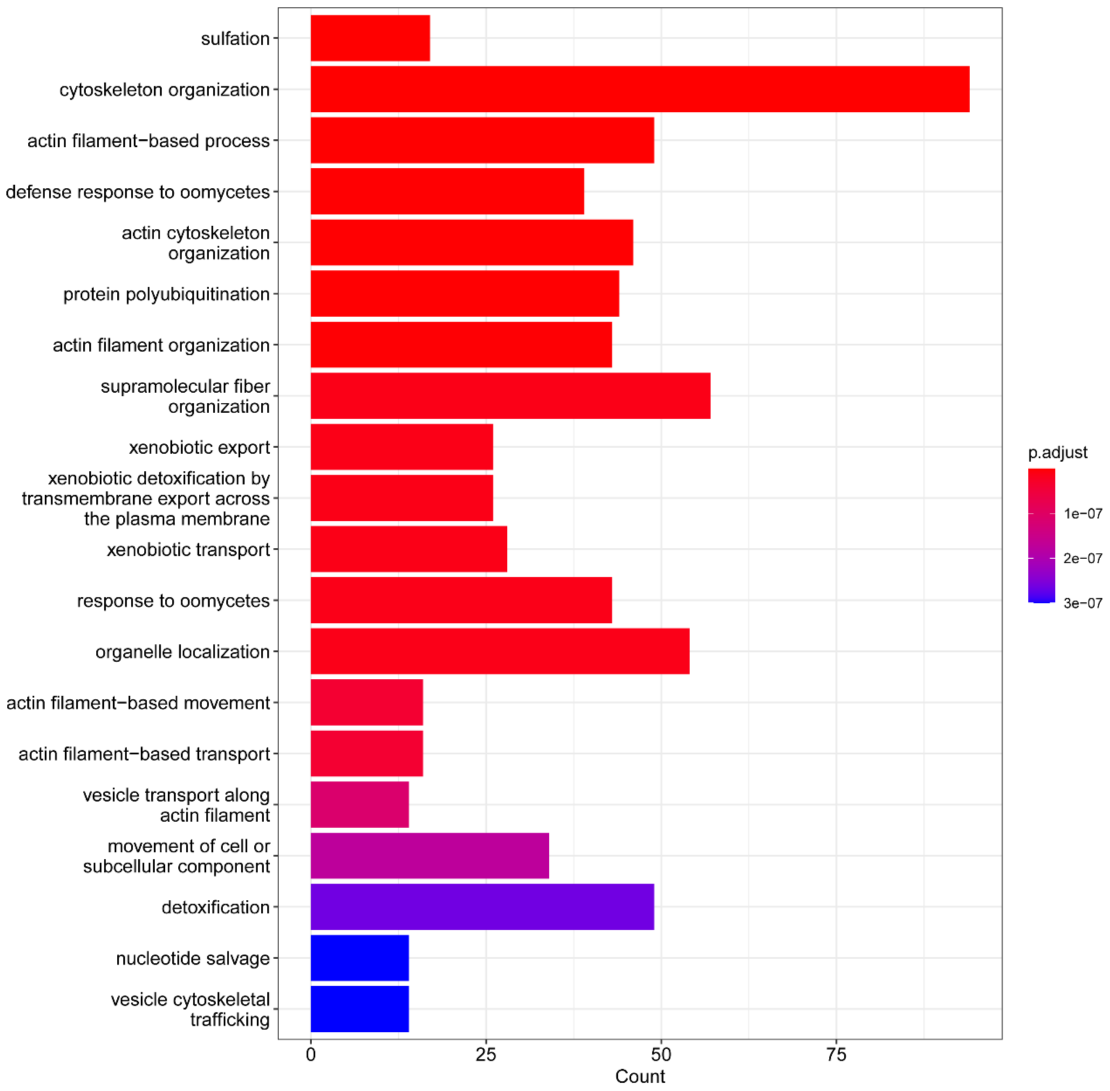


**Fig. S8. Top 20 enriched GO terms of convergently contracted gene families in alpine plant genomes.**

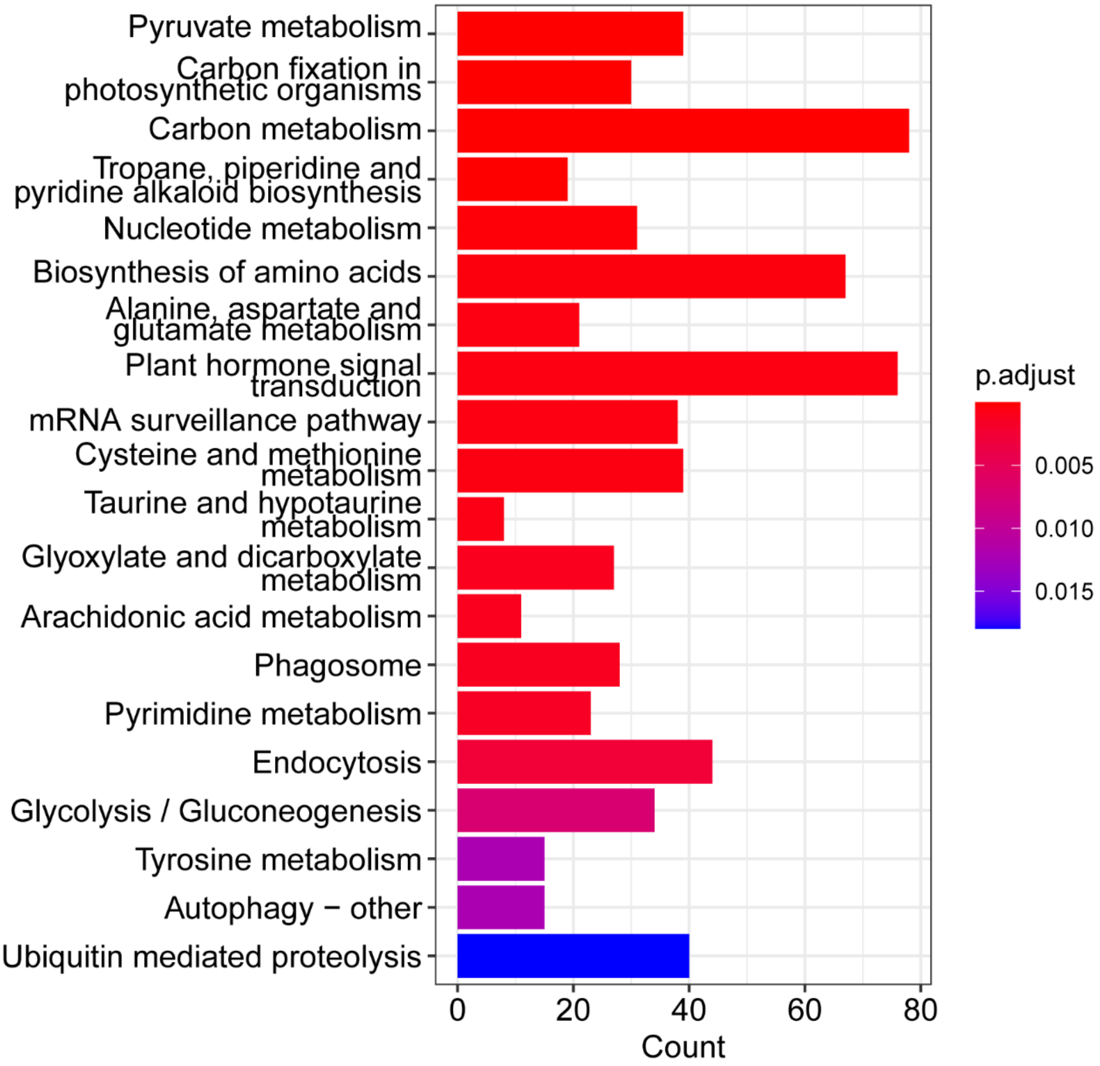


**Fig. S9. Top 20 enriched KEGG pathways of convergently contracted gene families in alpine plant genomes.**

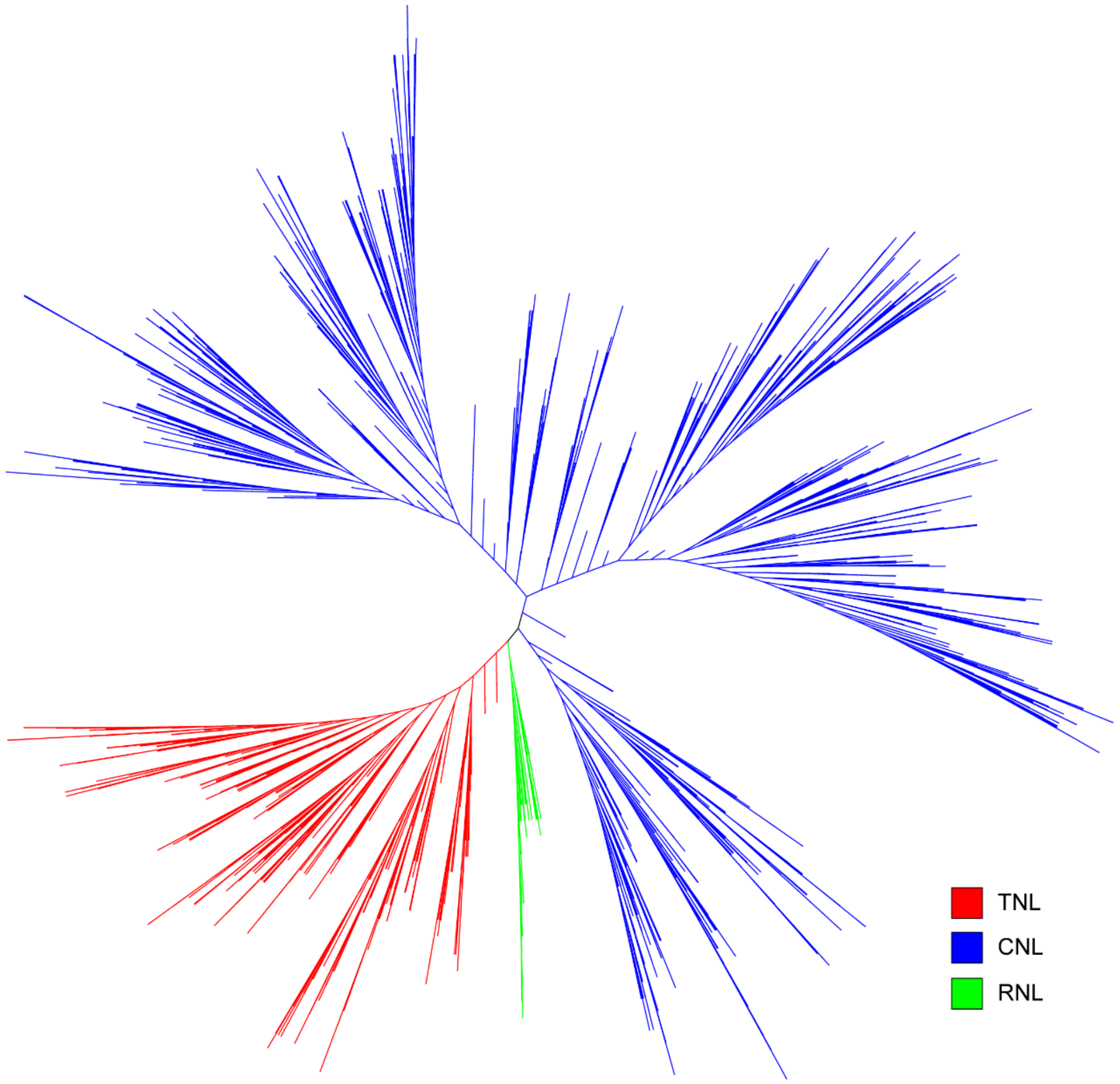


**Fig. S10. Gene tree of the nucleotide-binding site and leucine-rich-repeat domain receptor (NBS-LRR) genes.** Three subclasses, TIR-NBS-LRR (TNL), CC-NBS-LRR (CNL) and RPW8-NBS-LRR (RNL) with 1,024, 4,058, and 173 genes are included.

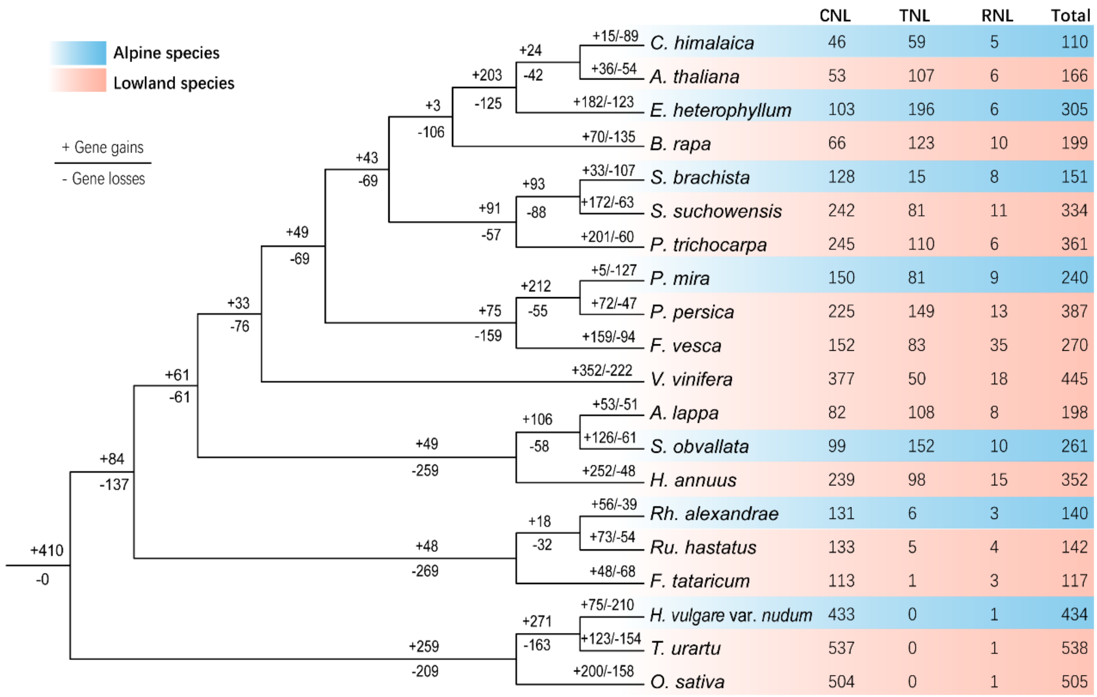


**Fig. S11. Loss and gain events of NBS-LRR genes in sampled species.** Three NBS-LRR gene subclasses, CC-NBS-LRR (CNL), TIR-NBS-LRR (TNL), and RPW8-NBS-LRR (RNL), are included. '-' or '+' indicates the number of gene losses and gains on each branch.

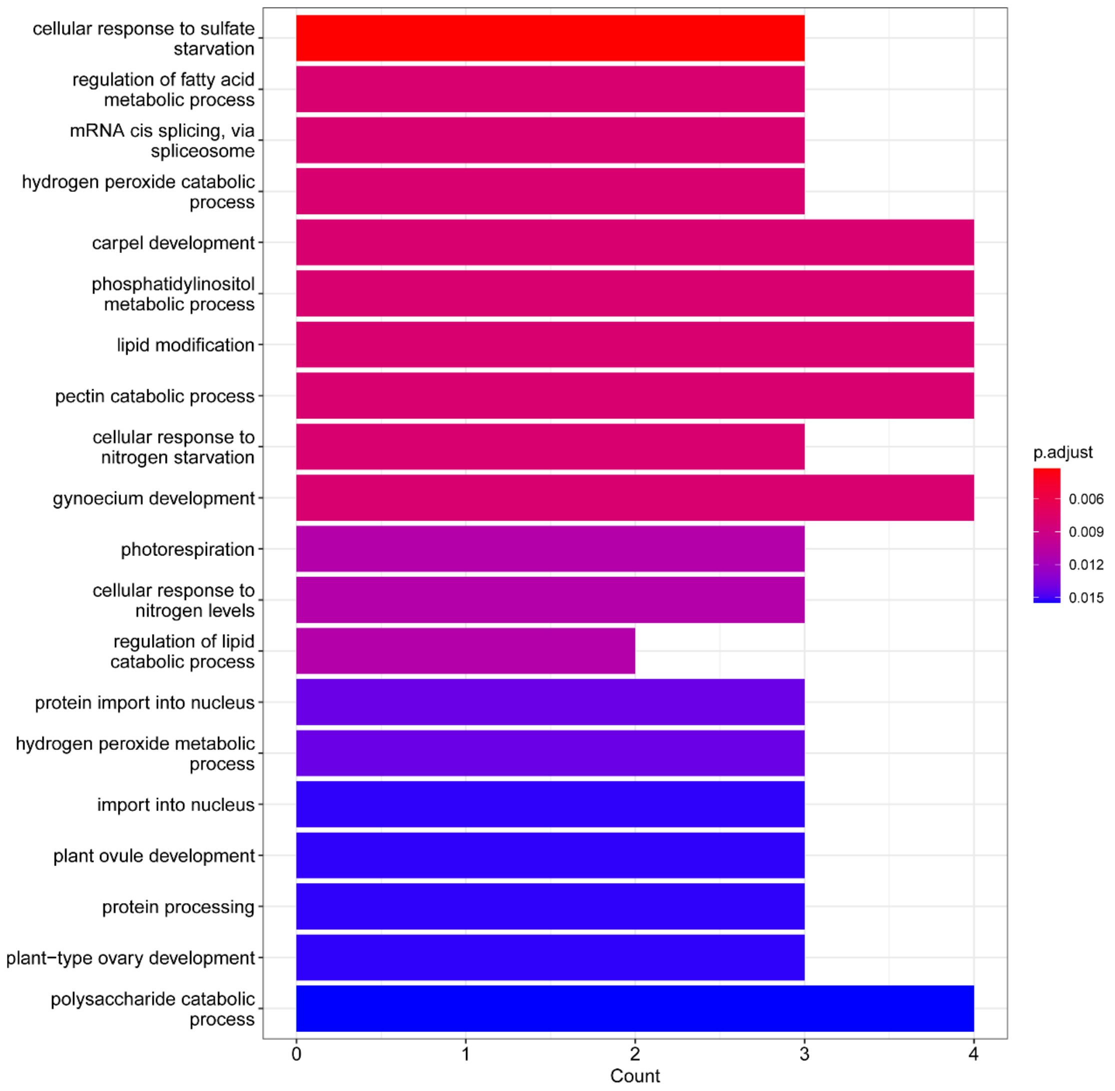


**Fig. S12. Top 20 enriched GO terms of genes undergoing convergent positive selection in alpine plant genomes.**

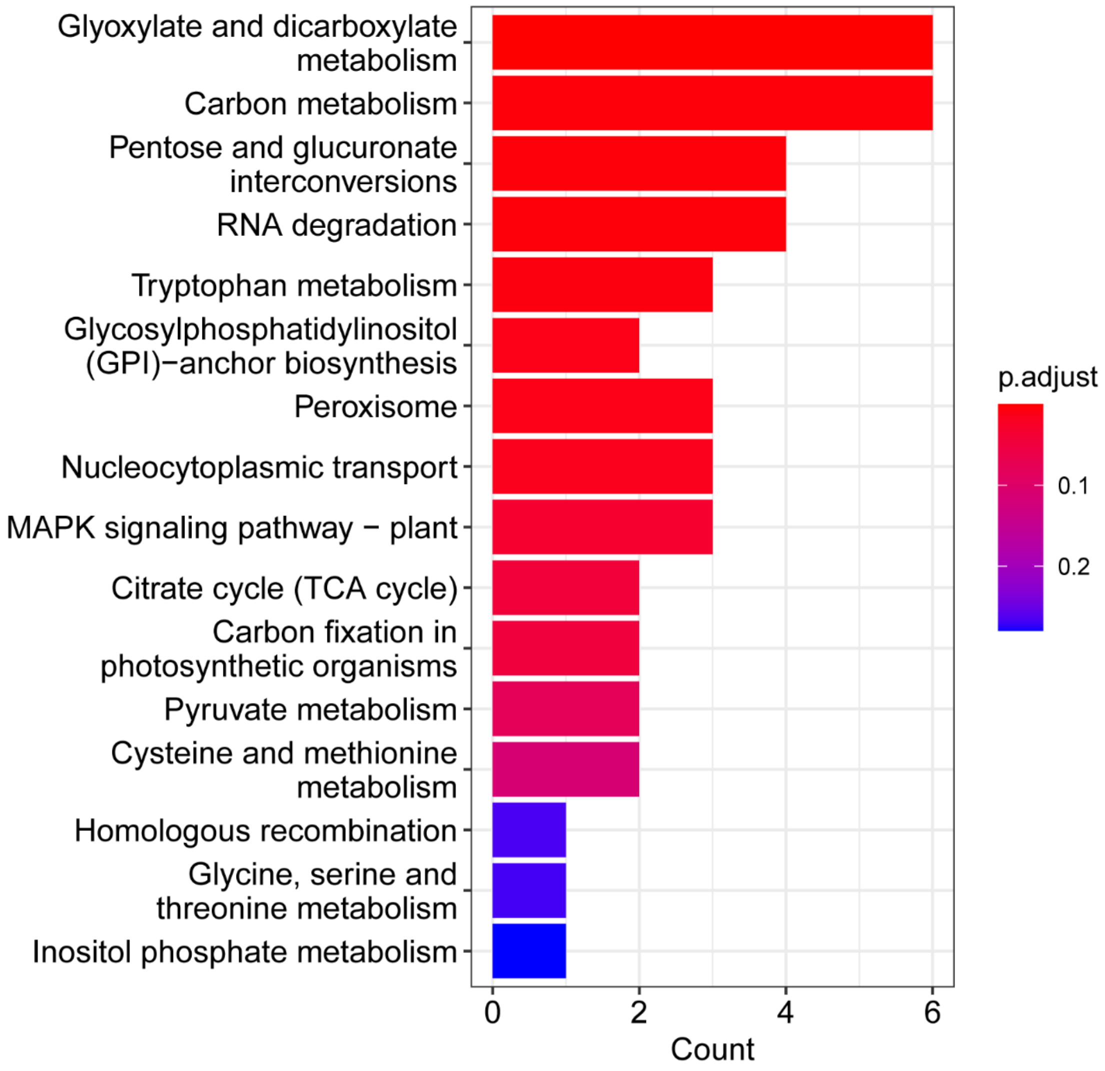


**Fig. S13. The enriched KEGG pathways of genes undergoing convergent positive selection in alpine plant genomes.**

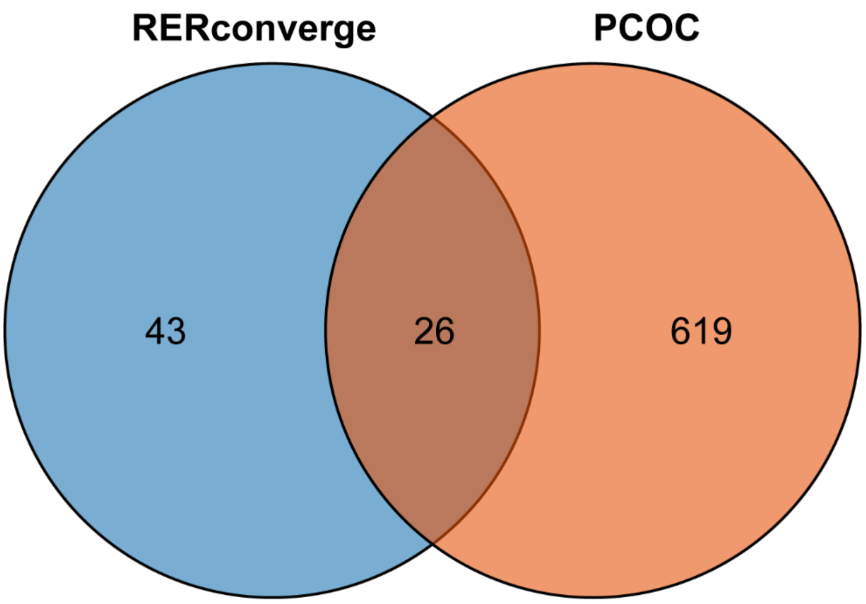


**Fig. S14. Veen plot showing the number of genes undergoing molecular convergence detected by RERconverge and PCOC analysis.**

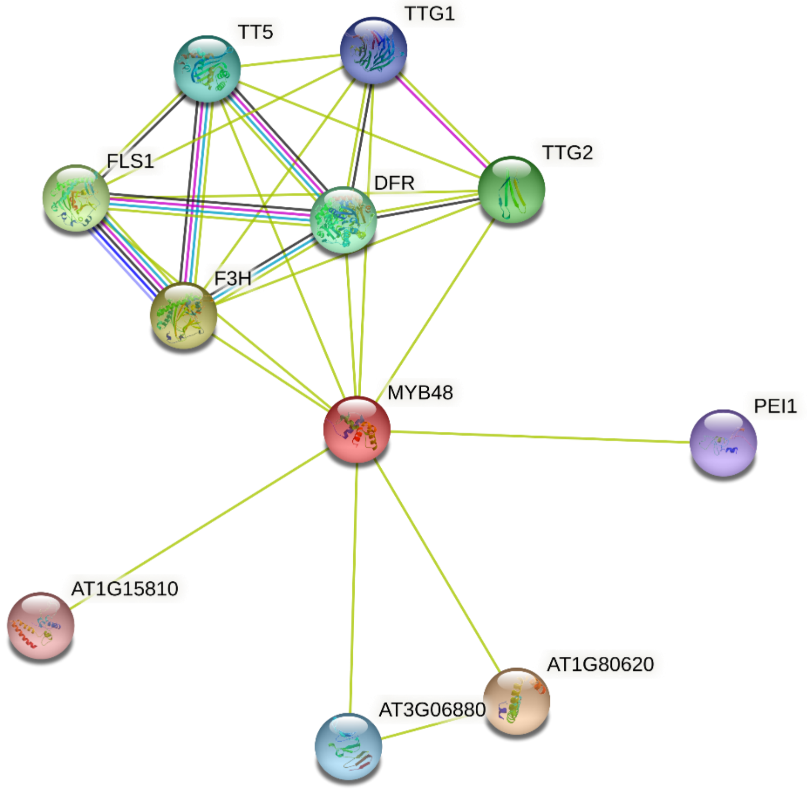


**Fig. S15. The predicted STRING network of Arabidopsis MYB48**. Network nodes represent proteins and edges represent protein-protein interactions.

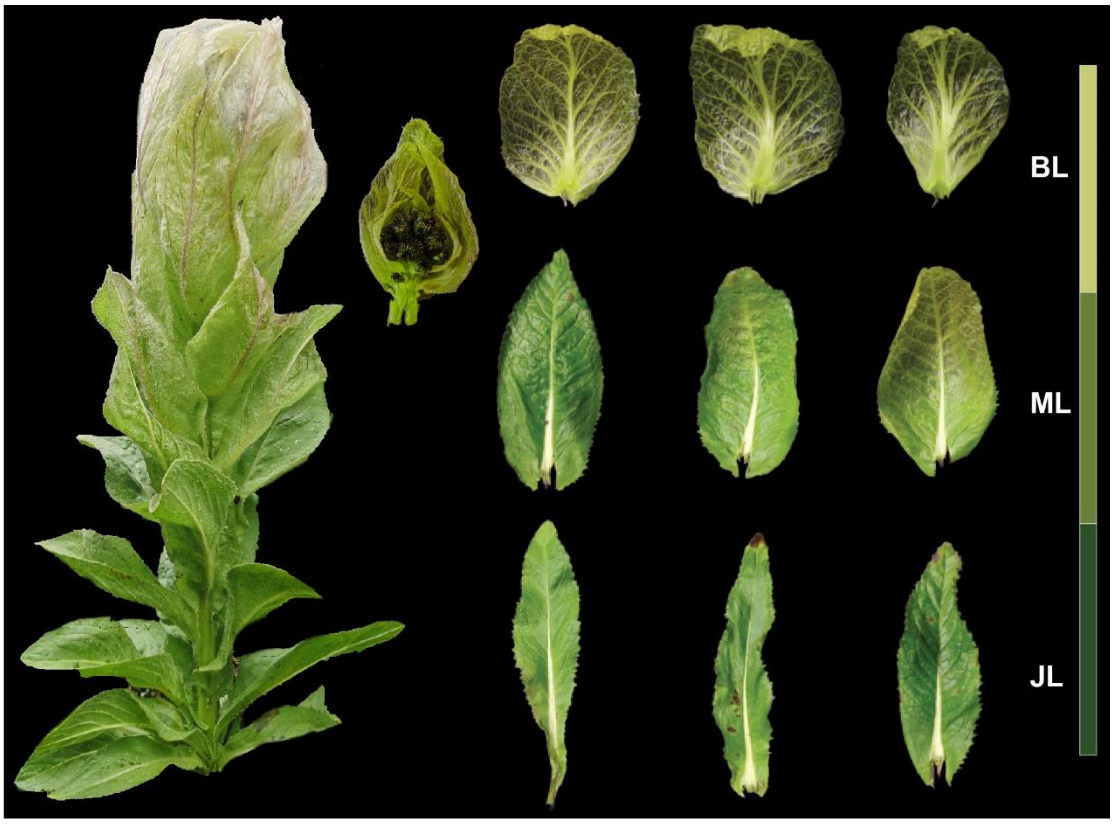


**Fig. S16. Sampling information of *S. obvallata* transcriptomic data.** Five tissues including three from leaves, basal leaves (JL), middle leaves (ML) and bract leaves (BL) as well as two from flowers and stems, were sampled.

**Table S1.** Genome sequences of 20 species used for comparative analysis, including seven alpine plants (bold) and 13 representative lowland sisters. NCBI: the National Center for Biotechnology Information. TAIR: The Arabidopsis Information Resource.

| Family | Species | Database |
|---|---|---|
| Brassicaceae | *A. thaliana* | TAIR |
| | *B. rapa* | Ensembl Plants |
| | ***C. himalaica*** | **NCBI** |
| | ***E. heterophyllum*** | **NCBI** |
| Polygonaceae | *F. tataricum* | Tartary Buckwheat Genome Project |
| | ***Rh. alexandrae*** | **Newly generated** |
| | *Ru. hastatus* | NCBI (manually annotated) |
| Rosaceae | *F. vesca* | Ensembl Plants |
| | ***P. mira*** | **Genome Database for Rosaceae** |
| | *P. persica* | Ensembl Plants |
| Asteraceae | *A. lappa* | NCBI |
| | *H. annuus* | Ensembl Plants |
| | ***S. obvallata*** | **Newly generated** |
| Poaceae | ***H. vulgare* var. *nudum*** | **Ensembl Plants** |
| | *O. sativa* | Ensembl Plants |
| | *T. urartu* | Ensembl Plants |
| Salicaceae | *P. trichocarpa* | Ensembl Plants |
| | ***S. brachista*** | **NCBI** |
| | *S. suchowensis* | NCBI |
| Vitaceae | *V. vinifera* | Ensembl Plants |

**Table S2.** Estimation of genome sizes for *S. obvallata* and *R. alexandrae* based on *K*-mer statistics.

| **Species** | ***K*-mer** | ***K*-mer count** | **Peak depth** | **Genome size (bp)** |
|---|---|---|---|---|
| *S. obvallata* | 17 | 78,809,693,159 | 34 | 2,250,587,609 |
| *R. alexandrae* | 17 | 92,775,947,880 | 42 | 2,084,855,059 |

**Table S3** The total sequencing data for genome assembly.

| **Species** | **Library** | **Total Base (bp)** | **Total Reads** | **Average Length (bp)** | **N50 (bp)** | **Q20 (%)** |
|---|---|---|---|---|---|---|
| ***S. obvallata*** | ONT | 143,303,536,196 | 11,784,529 | 22,917 | 60,615 | - |
| | Illumina | 95,142,395,400 | 634,282,636 | 150 | 150 | 96.11 |
| | Hi-C | 205,914,823,800 | 1,372,765,492 | 150 | 150 | 97.87 |
| ***R. alexandrae*** | ONT | 144,007,146,545 | 8,605,927 | 16,733 | 34,697 | - |
| | Illumina | 103,853,673,000 | 692,357,820 | 150, | 150 | 97.05 |

**Table S4** The information of transcriptomic data of *S. obvallata* used in the study.

| Tissue | Name | Total Base (bp) | Total Reads | Q20 (%) |
|---|---|---|---|---|
| Base leaf | JL-1 | 11,004,025,506 | 73,608,710 | 98.10 |
| | JL-2 | 10,494,272,308 | 70,156,674 | 98.26 |
| | JL-3 | 10,227,039,340 | 68,431,730 | 98.39 |
| Middle leaf | ML-1 | 10,422,933,558 | 69,743,294 | 98.38 |
| | ML-2 | 10,385,815,346 | 69,485,058 | 98.28 |
| | ML-3 | 10,277,264,344 | 68,692,336 | 98.28 |
| Bract leaf | BL-1 | 9,590,764,446 | 64,087,306 | 98.34 |
| | BL-2 | 10,989,175,450 | 73,458,712 | 98.30 |
| | BL-3 | 10,969,678,684 | 73,321,792 | 98.31 |
| Flower | F-1 | 11,435,046,272 | 76,473,416 | 98.27 |
| | F-2 | 10,349,894,180 | 69,276,056 | 98.19 |
| | F-3 | 10,134,792,130 | 67,828,610 | 98.24 |
| Stem | S-1 | 10,888,411,306 | 72,793,584 | 98.16 |
| | S-2 | 10,378,851,842 | 69,437,830 | 98.33 |
| | S-3 | 9,931,403,162 | 66,407,362 | 98.40 |
| Total | - | 157,479,367,874 | 1,053,202,470 | 98.28 |

**Table S5.** Statistics of initial genome assembly of *S. obvallata* and *R. alexandrae*.

| **Statistic** | ***S. obvallata*** | ***R. alexandrae*** |
|---|---|---|
| **Total length (bp)** | 2,044,030,733 | 2,039,881,226 |
| **Number of contigs** | 145 | 129 |
| **Largest contig (bp)** | 126,457,859 | 160,815,976 |
| **GC (%)** | 37.94 | 41.41 |
| **N50 (bp)** | 36,958,263 | 36,323,674 |
| **N75 (bp)** | 18,497,281 | 18,056,298 |

**Table S6.** Summary of the chromosome lengths of *S. obvallata* genome.

| Chromosome | Length (bp) |
|---|---:|
| chr1 | 91,194,000 |
| chr2 | 90,574,642 |
| chr3 | 82,964,075 |
| chr4 | 108,979,119 |
| chr5 | 201,268,011 |
| chr6 | 202,229,626 |
| chr7 | 87,271,487 |
| chr8 | 94,844,151 |
| chr9 | 173,197,562 |
| chr10 | 165,923,688 |
| chr11 | 125,735,984 |
| chr12 | 121,890,839 |
| chr13 | 120,237,955 |
| chr14 | 98,231,984 |
| chr15 | 107,747,241 |
| chr16 | 79,213,330 |
| Total | 1,951,503,694 |

**Table S7.** Genome completeness measured by Benchmarking Universal Single-Copy Orthologs (BUSCO) of *S. obvallata* genome.

| Type | Number | Percent (%) |
|---|---|---|
| **Complete BUSCOs (C)** | 1526 | 94.6 |
| Complete and single-copy BUSCOs (S) | 1420 | 88.0 |
| Complete and duplicated BUSCOs (D) | 106 | 6.6 |
| **Fragmented BUSCOs (F)** | 12 | 0.7 |
| **Missing BUSCOs (M)** | 76 | 4.7 |
| **Total BUSCO groups searched** | 1614 | 100 |

**Table S8.** Genome completeness measured by Benchmarking Universal Single-Copy Orthologs (BUSCO) of *R. alexandrae* genome.

| **Type** | **Number** | **Percent (%)** |
|---|---|---|
| **Complete BUSCOs (C)** | 2190 | 94.1 |
| Complete and single-copy BUSCOs (S) | 2017 | 86.7 |
| Complete and duplicated BUSCOs (D) | 173 | 7.4 |
| **Fragmented BUSCOs (F)** | 35 | 1.5 |
| **Missing BUSCOs (M)** | 101 | 4.4 |
| **Total BUSCO groups searched** | 2326 | 100 |

**Table S9.** Prediction of repetitive elements in the assembled *S. obvallata* genome.

| **TE type** | **Length (bp)** | **Percentage of sequence** |
|---|---:|---:|
| TE total | 1,597,806,979 | 81.88% |
| LINEs | 18,132,623 | 0.93% |
| LTR/Copia | 502,728,577 | 25.76% |
| LTR/Gypsy | 355,031,153 | 18.19% |
| DNA transposons | 39,617,754 | 2.03% |
| Unclassified | 657,623,138 | 33.70% |

**Table S10.** Prediction of repetitive elements in the assembled *R. alexandrae* genome.

| **TE type** | **Length (bp)** | **Percentage of sequence** |
|---|---:|---:|
| TE total | 1,665,563,021 | 81.65% |
| LINEs | 55,688,757 | 2.73% |
| LTR/Copia | 129,074,158 | 6.33% |
| LTR/Gypsy | 774,739,843 | 37.98% |
| DNA transposons | 832,271,540 | 4.08% |
| Unclassified | 592,993,472 | 29.07% |

**Table S11.** Functional annotation of predicted genes of *S. obvallata* and *R. alexandrae* genomes.

| **Database** | ***S. obvallata*** | | ***R. alexandrae*** | |
|---|---|---|---|---|
| | **Annotated proteins** | **Percent (%)** | **Annotated proteins** | **Percent (%)** |
| InterPro | 28,261 | 74.49 | 32,170 | 80.56 |
| NR | 36,433 | 96.03 | 47,268 | 76.90 |
| Swiss-Prot | 28,641 | 75.49 | 32,861 | 53.46 |
| eggNOG | 34,347 | 90.53 | 40,806 | 66.39 |
| Total annotated | 36,542 | 96.32 | 47,535 | 77.34 |
| Total genes | 37,938 | 100 | 61,463 | 100 |

**Table S12.** Summary of gene family clustering among the 20 species used.

| Species | Total Genes | Genes in families | Unassigned genes | Gene families | Unique families | Gene per family |
|---|---|---|---|---|---|---|
| *A. thaliana* | 48,455 | 46,894 | 1,561 | 15,545 | 571 | 3.7 |
| *A. lappa* | 32,771 | 30,564 | 2,207 | 14,663 | 402 | 4.5 |
| *B. rapa* | 41,025 | 38,865 | 2,160 | 14,968 | 558 | 5.4 |
| *C. himalaica* | 27,019 | 26,346 | 673 | 14,893 | 113 | 1.1 |
| *E. heterophyllum* | 42,586 | 40,279 | 2,307 | 15,341 | 646 | 5.6 |
| *F. tataricum* | 34,544 | 30,574 | 3,970 | 14,247 | 572 | 4.7 |
| *F. vesca* | 64,597 | 61,202 | 3,395 | 16,117 | 1,735 | 13.3 |
| *H. annuus* | 70,864 | 59,704 | 11,160 | 18,537 | 2,856 | 22.9 |
| *H. vulgare* var. *nudum* | 37,961 | 36,678 | 1,283 | 15,470 | 449 | 11.6 |
| *O. sativa* | 42,246 | 35,414 | 6,832 | 17,121 | 1,483 | 9.9 |
| *P. trichocarpa* | 73,012 | 68,079 | 4,933 | 16,535 | 930 | 4.0 |
| *P. mira* | 28,519 | 27,686 | 833 | 15,080 | 144 | 5.6 |
| *P. persica* | 47,089 | 45,081 | 2,008 | 15,438 | 363 | 2.6 |
| *R. alexandrae* | 61,463 | 57,793 | 3,670 | 17,798 | 2,670 | 27.2 |
| *R. hastatus* | 38,271 | 34,959 | 3,312 | 15,034 | 1,016 | 11.0 |
| *S. brachista* | 30,209 | 28,899 | 1,310 | 14,091 | 196 | 2.1 |
| *S. suchowensis* | 36,449 | 33,543 | 2,906 | 14,770 | 216 | 1.8 |
| *S. obvallata* | 37,938 | 36,338 | 1,600 | 15,095 | 503 | 4.6 |
| *T. urartu* | 41,449 | 34,952 | 6,497 | 16,308 | 1,335 | 9.4 |
| *V. vinifera* | 41,097 | 37,297 | 3,800 | 15,630 | 1,174 | 12.3 |

**Table S13.** Significantly enriched GO terms of convergently expanded gene families in alpine plant genomes based on the hypergeometric test. "*p*-adjust value" refers to the adjusted *p*-value of the Benjamini–Hochberg false discovery rate (FDR), and the same below.

| ID | Description | Gene Ratio | Background Ratio | *p*-adjust value |
|---|---|---|---|---|
| GO:0042542 | response to hydrogen peroxide | 10/156 | 76/26032 | 1.21E-08 |
| GO:0051259 | protein complex oligomerization | 9/156 | 56/26032 | 1.21E-08 |
| GO:0010817 | regulation of hormone levels | 13/156 | 303/26032 | 6.74E-06 |
| GO:0000302 | response to reactive oxygen species | 10/156 | 166/26032 | 8.60E-06 |
| GO:0010345 | suberin biosynthetic process | 5/156 | 22/26032 | 1.72E-05 |
| GO:0071456 | cellular response to hypoxia | 11/156 | 242/26032 | 1.72E-05 |
| GO:0036294 | cellular response to decreased oxygen levels | 11/156 | 244/26032 | 1.72E-05 |
| GO:0071453 | cellular response to oxygen levels | 11/156 | 244/26032 | 1.72E-05 |
| GO:0009150 | purine ribonucleotide metabolic process | 10/156 | 195/26032 | 1.72E-05 |
| GO:0006163 | purine nucleotide metabolic process | 10/156 | 203/26032 | 2.24E-05 |
| GO:0009117 | nucleotide metabolic process | 11/156 | 272/26032 | 3.05E-05 |
| GO:0006074 | (1->3)-beta-D-glucan metabolic process | 4/156 | 13/26032 | 3.05E-05 |
| GO:0006075 | (1->3)-beta-D-glucan biosynthetic process | 4/156 | 13/26032 | 3.05E-05 |
| GO:0036293 | response to decreased oxygen levels | 12/156 | 335/26032 | 3.05E-05 |
| GO:0070482 | response to oxygen levels | 12/156 | 336/26032 | 3.05E-05 |
| GO:0009259 | ribonucleotide metabolic process | 10/156 | 221/26032 | 3.05E-05 |
| GO:0006753 | nucleoside phosphate metabolic process | 11/156 | 282/26032 | 3.40E-05 |
| GO:0001709 | cell fate determination | 4/156 | 14/26032 | 3.40E-05 |
| GO:0072521 | purine-containing compound metabolic process | 10/156 | 228/26032 | 3.40E-05 |
| GO:0019693 | ribose phosphate metabolic process | 10/156 | 230/26032 | 3.50E-05 |
| GO:0009926 | auxin polar transport | 7/156 | 100/26032 | 5.98E-05 |
| GO:0006457 | protein folding | 9/156 | 200/26032 | 8.23E-05 |
| GO:0001666 | response to hypoxia | 11/156 | 327/26032 | 0.000108 |
| GO:0009851 | auxin biosynthetic process | 5/156 | 44/26032 | 0.00014 |
| GO:0006637 | acyl-CoA metabolic process | 5/156 | 49/26032 | 0.000208 |
| GO:0035383 | thioester metabolic process | 5/156 | 49/26032 | 0.000208 |
| GO:1901568 | fatty acid derivative metabolic process | 5/156 | 49/26032 | 0.000208 |
| GO:0060918 | auxin transport | 7/156 | 126/26032 | 0.000208 |
| GO:0009914 | hormone transport | 7/156 | 129/26032 | 0.000234 |
| GO:0009408 | response to heat | 10/156 | 299/26032 | 0.000237 |
| GO:0055086 | nucleobase-containing small molecule metabolic process | 11/156 | 380/26032 | 0.000325 |
| GO:0042446 | hormone biosynthetic process | 6/156 | 105/26032 | 0.000669 |
| GO:0033865 | nucleoside bisphosphate metabolic process | 5/156 | 66/26032 | 0.000711 |
| GO:0033875 | ribonucleoside bisphosphate metabolic process | 5/156 | 66/26032 | 0.000711 |
| GO:0034032 | purine nucleoside bisphosphate metabolic process | 5/156 | 66/26032 | 0.000711 |
| GO:0009850 | auxin metabolic process | 5/156 | 75/26032 | 0.00127 |

| | | | | |
|---|---|---|---|---|
| GO:0010540 | basipetal auxin transport | 3/156 | 15/26032 | 0.00127 |
| GO:0045165 | cell fate commitment | 5/156 | 100/26032 | 0.004686 |
| GO:0042445 | hormone metabolic process | 6/156 | 160/26032 | 0.005523 |
| GO:0009142 | nucleoside triphosphate biosynthetic process | 4/156 | 60/26032 | 0.006013 |
| GO:0010150 | leaf senescence | 7/156 | 230/26032 | 0.00617 |
| GO:0009699 | phenylpropanoid biosynthetic process | 5/156 | 111/26032 | 0.006847 |
| GO:0009638 | phototropism | 3/156 | 28/26032 | 0.007415 |
| GO:0009141 | nucleoside triphosphate metabolic process | 4/156 | 65/26032 | 0.007415 |
| GO:0090693 | plant organ senescence | 7/156 | 248/26032 | 0.008773 |
| GO:0016998 | cell wall macromolecule catabolic process | 3/156 | 33/26032 | 0.01131 |
| GO:0009723 | response to ethylene | 7/156 | 261/26032 | 0.01131 |
| GO:0006754 | ATP biosynthetic process | 3/156 | 34/26032 | 0.011607 |
| GO:0015986 | proton motive force-driven ATP synthesis | 3/156 | 34/26032 | 0.011607 |
| GO:0044264 | cellular polysaccharide metabolic process | 8/156 | 353/26032 | 0.013909 |
| GO:0051274 | beta-glucan biosynthetic process | 4/156 | 82/26032 | 0.015277 |
| GO:0009698 | phenylpropanoid metabolic process | 5/156 | 145/26032 | 0.018099 |
| GO:0006545 | glycine biosynthetic process | 2/156 | 11/26032 | 0.018099 |
| GO:0080086 | stamen filament development | 2/156 | 11/26032 | 0.018099 |
| GO:0051273 | beta-glucan metabolic process | 4/156 | 89/26032 | 0.018483 |
| GO:0009205 | purine ribonucleoside triphosphate metabolic process | 3/156 | 42/26032 | 0.018483 |
| GO:0009206 | purine ribonucleoside triphosphate biosynthetic process | 3/156 | 42/26032 | 0.018483 |
| GO:0046034 | ATP metabolic process | 5/156 | 150/26032 | 0.018998 |
| GO:0009145 | purine nucleoside triphosphate biosynthetic process | 3/156 | 43/26032 | 0.019112 |
| GO:0044036 | cell wall macromolecule metabolic process | 6/156 | 228/26032 | 0.022181 |
| GO:0009144 | purine nucleoside triphosphate metabolic process | 3/156 | 46/26032 | 0.022181 |
| GO:0006566 | threonine metabolic process | 2/156 | 13/26032 | 0.022181 |
| GO:0009637 | response to blue light | 5/156 | 163/26032 | 0.025047 |
| GO:0009733 | response to auxin | 8/156 | 414/26032 | 0.029032 |
| GO:0009199 | ribonucleoside triphosphate metabolic process | 3/156 | 54/26032 | 0.032729 |
| GO:0009201 | ribonucleoside triphosphate biosynthetic process | 3/156 | 54/26032 | 0.032729 |
| GO:0006544 | glycine metabolic process | 2/156 | 18/26032 | 0.039478 |
| GO:0006119 | oxidative phosphorylation | 3/156 | 63/26032 | 0.048904 |

**Table S14.** Significantly enriched KEGG pathways of convergently expanded gene families in alpine plant genomes based on the hypergeometric test.

| **ID** | **Description** | **Gene Ratio** | **Background Ratio** | ***p*-adjust value** |
|---|---|---|---|---|
| ath00073 | Cutin, suberine and wax biosynthesis | 5/53 | 37/5098 | 0.000883 |
| ath00190 | Oxidative phosphorylation | 8/53 | 163/5098 | 0.002715 |
| ath04141 | Protein processing in endoplasmic reticulum | 9/53 | 215/5098 | 0.002715 |
| ath00380 | Tryptophan metabolism | 5/53 | 64/5098 | 0.003121 |
| ath04146 | Peroxisome | 5/53 | 87/5098 | 0.010104 |
| ath00260 | Glycine, serine and threonine metabolism | 4/53 | 70/5098 | 0.02493 |

**Table S15.** Significantly enriched GO terms of convergently contracted gene families in alpine plant genomes based on the hypergeometric test. Only p-adjust value < 0.01are provided.

| ID | Description | Gene Ratio | Background Ratio | *p*-adjust value |
|---|---|---|---|---|
| GO:0051923 | sulfation | 17/3445 | 17/26032 | 2.28E-12 |
| GO:0007010 | cytoskeleton organization | 94/3445 | 325/26032 | 6.17E-11 |
| GO:0030029 | actin filament-based process | 49/3445 | 128/26032 | 6.52E-10 |
| GO:0002229 | defense response to oomycetes | 39/3445 | 91/26032 | 1.14E-09 |
| GO:0030036 | actin cytoskeleton organization | 46/3445 | 119/26032 | 1.14E-09 |
| GO:0000209 | protein polyubiquitination | 44/3445 | 111/26032 | 1.14E-09 |
| GO:0007015 | actin filament organization | 43/3445 | 109/26032 | 2.08E-09 |
| GO:0097435 | supramolecular fiber organization | 57/3445 | 177/26032 | 1.26E-08 |
| GO:0046618 | xenobiotic export | 26/3445 | 50/26032 | 1.26E-08 |
| GO:1990961 | xenobiotic detoxification by transmembrane export across the plasma membrane | 26/3445 | 50/26032 | 1.26E-08 |
| GO:0042908 | xenobiotic transport | 28/3445 | 57/26032 | 1.26E-08 |
| GO:0002239 | response to oomycetes | 43/3445 | 116/26032 | 1.34E-08 |
| GO:0051640 | organelle localization | 54/3445 | 165/26032 | 1.34E-08 |
| GO:0030048 | actin filament-based movement | 16/3445 | 22/26032 | 3.89E-08 |
| GO:0099515 | actin filament-based transport | 16/3445 | 22/26032 | 3.89E-08 |
| GO:0030050 | vesicle transport along actin filament | 14/3445 | 18/26032 | 1.13E-07 |
| GO:0006928 | movement of cell or subcellular component | 34/3445 | 87/26032 | 1.75E-07 |
| GO:0098754 | detoxification | 49/3445 | 155/26032 | 2.64E-07 |
| GO:0043173 | nucleotide salvage | 14/3445 | 19/26032 | 3.00E-07 |
| GO:0099518 | vesicle cytoskeletal trafficking | 14/3445 | 19/26032 | 3.00E-07 |
| GO:0051648 | vesicle localization | 21/3445 | 40/26032 | 3.22E-07 |
| GO:0051650 | establishment of vesicle localization | 21/3445 | 40/26032 | 3.22E-07 |
| GO:0140115 | export across plasma membrane | 34/3445 | 91/26032 | 5.09E-07 |
| GO:0043547 | positive regulation of GTPase activity | 20/3445 | 40/26032 | 2.07E-06 |
| GO:0009896 | positive regulation of catabolic process | 43/3445 | 138/26032 | 2.84E-06 |
| GO:0015914 | phospholipid transport | 13/3445 | 19/26032 | 3.60E-06 |
| GO:0009636 | response to toxic substance | 51/3445 | 180/26032 | 4.74E-06 |
| GO:0140352 | export from cell | 50/3445 | 175/26032 | 4.74E-06 |
| GO:0043087 | regulation of GTPase activity | 20/3445 | 43/26032 | 8.02E-06 |
| GO:0022406 | membrane docking | 22/3445 | 51/26032 | 9.34E-06 |
| GO:0140056 | organelle localization by membrane tethering | 22/3445 | 51/26032 | 9.34E-06 |
| GO:0031331 | positive regulation of cellular catabolic process | 40/3445 | 130/26032 | 9.58E-06 |
| GO:0030705 | cytoskeleton-dependent intracellular transport | 17/3445 | 33/26032 | 9.69E-06 |
| GO:0043094 | cellular metabolic compound salvage | 29/3445 | 81/26032 | 1.30E-05 |
| GO:0006325 | chromatin organization | 66/3445 | 268/26032 | 1.79E-05 |
| GO:0051345 | positive regulation of hydrolase activity | 20/3445 | 46/26032 | 2.53E-05 |
| GO:0031497 | chromatin assembly | 35/3445 | 112/26032 | 3.21E-05 |

| | | | | |
|---|---|---|---|---|
| GO:0009845 | seed germination | 49/3445 | 184/26032 | 4.78E-05 |
| GO:0032436 | positive regulation of proteasomal ubiquitin-dependent protein catabolic process | 21/3445 | 53/26032 | 7.72E-05 |
| GO:0009123 | nucleoside monophosphate metabolic process | 20/3445 | 49/26032 | 7.72E-05 |
| GO:0051656 | establishment of organelle localization | 32/3445 | 103/26032 | 9.98E-05 |
| GO:0006334 | nucleosome assembly | 18/3445 | 42/26032 | 0.000105 |
| GO:0045732 | positive regulation of protein catabolic process | 24/3445 | 67/26032 | 0.00011 |
| GO:0030243 | cellulose metabolic process | 26/3445 | 76/26032 | 0.000113 |
| GO:0051247 | positive regulation of protein metabolic process | 41/3445 | 149/26032 | 0.000122 |
| GO:0006108 | malate metabolic process | 11/3445 | 18/26032 | 0.000122 |
| GO:0032434 | regulation of proteasomal ubiquitin-dependent protein catabolic process | 21/3445 | 55/26032 | 0.000126 |
| GO:1901800 | positive regulation of proteasomal protein catabolic process | 21/3445 | 55/26032 | 0.000126 |
| GO:2000060 | positive regulation of ubiquitin-dependent protein catabolic process | 21/3445 | 55/26032 | 0.000126 |
| GO:0065004 | protein-DNA complex assembly | 28/3445 | 86/26032 | 0.000126 |
| GO:0009124 | nucleoside monophosphate biosynthetic process | 19/3445 | 47/26032 | 0.000131 |
| GO:0071215 | cellular response to abscisic acid stimulus | 75/3445 | 338/26032 | 0.000143 |
| GO:0097306 | cellular response to alcohol | 75/3445 | 338/26032 | 0.000143 |
| GO:0046777 | protein autophosphorylation | 46/3445 | 177/26032 | 0.000147 |
| GO:0046112 | nucleobase biosynthetic process | 13/3445 | 25/26032 | 0.000149 |
| GO:0009966 | regulation of signal transduction | 89/3445 | 421/26032 | 0.000149 |
| GO:0018958 | phenol-containing compound metabolic process | 24/3445 | 69/26032 | 0.000151 |
| GO:0034728 | nucleosome organization | 20/3445 | 52/26032 | 0.000159 |
| GO:1901605 | alpha-amino acid metabolic process | 81/3445 | 375/26032 | 0.000161 |
| GO:0006869 | lipid transport | 38/3445 | 137/26032 | 0.000173 |
| GO:0045332 | phospholipid translocation | 9/3445 | 13/26032 | 0.000176 |
| GO:0032270 | positive regulation of cellular protein metabolic process | 38/3445 | 138/26032 | 0.000197 |
| GO:0006220 | pyrimidine nucleotide metabolic process | 15/3445 | 33/26032 | 0.000197 |
| GO:0006221 | pyrimidine nucleotide biosynthetic process | 15/3445 | 33/26032 | 0.000197 |
| GO:0006305 | DNA alkylation | 20/3445 | 53/26032 | 0.000197 |
| GO:0006306 | DNA methylation | 20/3445 | 53/26032 | 0.000197 |
| GO:0043648 | dicarboxylic acid metabolic process | 28/3445 | 89/26032 | 0.000199 |
| GO:0006338 | chromatin remodeling | 30/3445 | 99/26032 | 0.000214 |
| GO:0071824 | protein-DNA complex subunit organization | 30/3445 | 99/26032 | 0.000214 |
| GO:1903052 | positive regulation of proteolysis involved in cellular protein catabolic process | 21/3445 | 58/26032 | 0.000232 |
| GO:2000058 | regulation of ubiquitin-dependent protein catabolic process | 21/3445 | 58/26032 | 0.000232 |
| GO:0009132 | nucleoside diphosphate metabolic process | 25/3445 | 76/26032 | 0.000233 |
| GO:0042537 | benzene-containing compound metabolic process | 25/3445 | 76/26032 | 0.000233 |
| GO:0008154 | actin polymerization or depolymerization | 18/3445 | 46/26032 | 0.000278 |
| GO:0000281 | mitotic cytokinesis | 17/3445 | 42/26032 | 0.000278 |

| | | | | |
|---|---|---|---|---|
| GO:0006538 | glutamate catabolic process | 8/3445 | 11/26032 | 0.000278 |
| GO:0006904 | vesicle docking involved in exocytosis | 8/3445 | 11/26032 | 0.000278 |
| GO:0023051 | regulation of signaling | 89/3445 | 431/26032 | 0.000279 |
| GO:0045862 | positive regulation of proteolysis | 21/3445 | 59/26032 | 0.000279 |
| GO:1903364 | positive regulation of cellular protein catabolic process | 21/3445 | 59/26032 | 0.000279 |
| GO:0009894 | regulation of catabolic process | 49/3445 | 200/26032 | 0.000282 |
| GO:0006536 | glutamate metabolic process | 13/3445 | 27/26032 | 0.000299 |
| GO:0009738 | abscisic acid-activated signaling pathway | 61/3445 | 268/26032 | 0.00031 |
| GO:0034204 | lipid translocation | 9/3445 | 14/26032 | 0.000312 |
| GO:0090158 | endoplasmic reticulum membrane organization | 9/3445 | 14/26032 | 0.000312 |
| GO:0030244 | cellulose biosynthetic process | 23/3445 | 69/26032 | 0.000351 |
| GO:0072528 | pyrimidine-containing compound biosynthetic process | 17/3445 | 43/26032 | 0.000356 |
| GO:0032268 | regulation of cellular protein metabolic process | 82/3445 | 393/26032 | 0.000367 |
| GO:0010646 | regulation of cell communication | 90/3445 | 442/26032 | 0.000389 |
| GO:0061640 | cytoskeleton-dependent cytokinesis | 22/3445 | 65/26032 | 0.000389 |
| GO:0048584 | positive regulation of response to stimulus | 58/3445 | 254/26032 | 0.000411 |
| GO:0009696 | salicylic acid metabolic process | 19/3445 | 52/26032 | 0.000411 |
| GO:0051246 | regulation of protein metabolic process | 86/3445 | 419/26032 | 0.000418 |
| GO:0009833 | plant-type primary cell wall biogenesis | 13/3445 | 28/26032 | 0.000422 |
| GO:0009185 | ribonucleoside diphosphate metabolic process | 21/3445 | 61/26032 | 0.000422 |
| GO:0051273 | beta-glucan metabolic process | 27/3445 | 89/26032 | 0.000422 |
| GO:0009161 | ribonucleoside monophosphate metabolic process | 17/3445 | 44/26032 | 0.000457 |
| GO:0045893 | positive regulation of transcription, DNA-templated | 99/3445 | 500/26032 | 0.00046 |
| GO:0042026 | protein refolding | 15/3445 | 36/26032 | 0.00047 |
| GO:0090351 | seedling development | 50/3445 | 211/26032 | 0.000489 |
| GO:0009789 | positive regulation of abscisic acid-activated signaling pathway | 19/3445 | 53/26032 | 0.000494 |
| GO:0006241 | CTP biosynthetic process | 10/3445 | 18/26032 | 0.000494 |
| GO:0009208 | pyrimidine ribonucleoside triphosphate metabolic process | 10/3445 | 18/26032 | 0.000494 |
| GO:0009209 | pyrimidine ribonucleoside triphosphate biosynthetic process | 10/3445 | 18/26032 | 0.000494 |
| GO:0046036 | CTP metabolic process | 10/3445 | 18/26032 | 0.000494 |
| GO:0061136 | regulation of proteasomal protein catabolic process | 21/3445 | 62/26032 | 0.000503 |
| GO:0030388 | fructose 1,6-bisphosphate metabolic process | 8/3445 | 12/26032 | 0.000524 |
| GO:0046033 | AMP metabolic process | 8/3445 | 12/26032 | 0.000524 |
| GO:0009299 | mRNA transcription | 9/3445 | 15/26032 | 0.000536 |
| GO:0009593 | detection of chemical stimulus | 13/3445 | 29/26032 | 0.000583 |
| GO:0000097 | sulfur amino acid biosynthetic process | 16/3445 | 41/26032 | 0.000599 |
| GO:0006096 | glycolytic process | 19/3445 | 54/26032 | 0.000609 |
| GO:0006757 | ATP generation from ADP | 19/3445 | 54/26032 | 0.000609 |
| GO:0008643 | carbohydrate transport | 21/3445 | 63/26032 | 0.000609 |
| GO:0051606 | detection of stimulus | 21/3445 | 63/26032 | 0.000609 |
| GO:0071496 | cellular response to external stimulus | 56/3445 | 249/26032 | 0.000698 |

| | | | | |
|---|---|---|---|---|
| GO:0031329 | regulation of cellular catabolic process | 43/3445 | 176/26032 | 0.000698 |
| GO:0030042 | actin filament depolymerization | 12/3445 | 26/26032 | 0.000768 |
| GO:0050801 | ion homeostasis | 79/3445 | 386/26032 | 0.000772 |
| GO:0009156 | ribonucleoside monophosphate biosynthetic process | 16/3445 | 42/26032 | 0.000786 |
| GO:0009148 | pyrimidine nucleoside triphosphate biosynthetic process | 10/3445 | 19/26032 | 0.000795 |
| GO:2000026 | regulation of multicellular organismal development | 97/3445 | 497/26032 | 0.000795 |
| GO:0048278 | vesicle docking | 15/3445 | 38/26032 | 0.000811 |
| GO:0006167 | AMP biosynthetic process | 7/3445 | 10/26032 | 0.000925 |
| GO:0008655 | pyrimidine-containing compound salvage | 7/3445 | 10/26032 | 0.000925 |
| GO:0043102 | amino acid salvage | 7/3445 | 10/26032 | 0.000925 |
| GO:0045596 | negative regulation of cell differentiation | 7/3445 | 10/26032 | 0.000925 |
| GO:0071267 | L-methionine salvage | 7/3445 | 10/26032 | 0.000925 |
| GO:0009135 | purine nucleoside diphosphate metabolic process | 19/3445 | 56/26032 | 0.000947 |
| GO:0009179 | purine ribonucleoside diphosphate metabolic process | 19/3445 | 56/26032 | 0.000947 |
| GO:0046031 | ADP metabolic process | 19/3445 | 56/26032 | 0.000947 |
| GO:0030001 | metal ion transport | 51/3445 | 224/26032 | 0.000952 |
| GO:0030837 | negative regulation of actin filament polymerization | 8/3445 | 13/26032 | 0.000971 |
| GO:0140029 | exocytic process | 8/3445 | 13/26032 | 0.000971 |
| GO:0044728 | DNA methylation or demethylation | 23/3445 | 75/26032 | 0.000998 |
| GO:0016311 | dephosphorylation | 50/3445 | 219/26032 | 0.001005 |
| GO:0048580 | regulation of post-embryonic development | 84/3445 | 421/26032 | 0.001005 |
| GO:0048467 | gynoecium development | 28/3445 | 100/26032 | 0.001038 |
| GO:0034220 | ion transmembrane transport | 67/3445 | 319/26032 | 0.001073 |
| GO:1903050 | regulation of proteolysis involved in cellular protein catabolic process | 21/3445 | 66/26032 | 0.001086 |
| GO:0009147 | pyrimidine nucleoside triphosphate metabolic process | 10/3445 | 20/26032 | 0.001207 |
| GO:0009826 | unidimensional cell growth | 85/3445 | 431/26032 | 0.00134 |
| GO:0002221 | pattern recognition receptor signaling pathway | 11/3445 | 24/26032 | 0.001431 |
| GO:0051274 | beta-glucan biosynthetic process | 24/3445 | 82/26032 | 0.001493 |
| GO:0010629 | negative regulation of gene expression | 87/3445 | 445/26032 | 0.001493 |
| GO:0009129 | pyrimidine nucleoside monophosphate metabolic process | 9/3445 | 17/26032 | 0.001493 |
| GO:0009218 | pyrimidine ribonucleotide metabolic process | 12/3445 | 28/26032 | 0.001493 |
| GO:0009220 | pyrimidine ribonucleotide biosynthetic process | 12/3445 | 28/26032 | 0.001493 |
| GO:0051094 | positive regulation of developmental process | 35/3445 | 140/26032 | 0.001679 |
| GO:0048582 | positive regulation of post-embryonic development | 25/3445 | 88/26032 | 0.001765 |
| GO:0019856 | pyrimidine nucleobase biosynthetic process | 8/3445 | 14/26032 | 0.001765 |
| GO:0043649 | dicarboxylic acid catabolic process | 8/3445 | 14/26032 | 0.001765 |
| GO:0070262 | peptidyl-serine dephosphorylation | 7/3445 | 11/26032 | 0.001872 |
| GO:0071265 | L-methionine biosynthetic process | 7/3445 | 11/26032 | 0.001872 |
| GO:0006470 | protein dephosphorylation | 35/3445 | 141/26032 | 0.001877 |
| GO:0031668 | cellular response to extracellular stimulus | 53/3445 | 243/26032 | 0.001912 |
| GO:1903362 | regulation of cellular protein catabolic process | 21/3445 | 69/26032 | 0.001975 |
| GO:0006098 | pentose-phosphate shunt | 13/3445 | 33/26032 | 0.002035 |
| GO:0009991 | response to extracellular stimulus | 64/3445 | 309/26032 | 0.002035 |

| | | | | |
|---|---|---|---|---|
| GO:0006090 | pyruvate metabolic process | 24/3445 | 84/26032 | 0.002035 |
| GO:0015748 | organophosphate ester transport | 16/3445 | 46/26032 | 0.002071 |
| GO:0007018 | microtubule-based movement | 20/3445 | 65/26032 | 0.002295 |
| GO:0009064 | glutamine family amino acid metabolic process | 20/3445 | 65/26032 | 0.002295 |
| GO:0097035 | regulation of membrane lipid distribution | 9/3445 | 18/26032 | 0.002377 |
| GO:0005984 | disaccharide metabolic process | 22/3445 | 75/26032 | 0.00239 |
| GO:0042176 | regulation of protein catabolic process | 24/3445 | 85/26032 | 0.00239 |
| GO:0006304 | DNA modification | 23/3445 | 80/26032 | 0.00239 |
| GO:0002764 | immune response-regulating signaling pathway | 14/3445 | 38/26032 | 0.002515 |
| GO:0005992 | trehalose biosynthetic process | 10/3445 | 22/26032 | 0.002722 |
| GO:0009395 | phospholipid catabolic process | 10/3445 | 22/26032 | 0.002722 |
| GO:0016052 | carbohydrate catabolic process | 50/3445 | 230/26032 | 0.002839 |
| GO:0032507 | maintenance of protein location in cell | 11/3445 | 26/26032 | 0.002839 |
| GO:0045185 | maintenance of protein location | 11/3445 | 26/26032 | 0.002839 |
| GO:0006535 | cysteine biosynthetic process from serine | 8/3445 | 15/26032 | 0.00287 |
| GO:0031333 | negative regulation of protein-containing complex assembly | 8/3445 | 15/26032 | 0.00287 |
| GO:0032272 | negative regulation of protein polymerization | 8/3445 | 15/26032 | 0.00287 |
| GO:0032776 | DNA methylation on cytosine | 8/3445 | 15/26032 | 0.00287 |
| GO:0098660 | inorganic ion transmembrane transport | 51/3445 | 237/26032 | 0.003112 |
| GO:0006165 | nucleoside diphosphate phosphorylation | 19/3445 | 62/26032 | 0.00318 |
| GO:0009311 | oligosaccharide metabolic process | 24/3445 | 87/26032 | 0.003245 |
| GO:0009733 | response to auxin | 80/3445 | 414/26032 | 0.003251 |
| GO:0006607 | NLS-bearing protein import into nucleus | 7/3445 | 12/26032 | 0.003329 |
| GO:0051643 | endoplasmic reticulum localization | 7/3445 | 12/26032 | 0.003329 |
| GO:0061817 | endoplasmic reticulum-plasma membrane tethering | 7/3445 | 12/26032 | 0.003329 |
| GO:0007017 | microtubule-based process | 52/3445 | 244/26032 | 0.003361 |
| GO:0048468 | cell development | 88/3445 | 465/26032 | 0.003361 |
| GO:0010214 | seed coat development | 17/3445 | 53/26032 | 0.003429 |
| GO:0006740 | NADPH regeneration | 13/3445 | 35/26032 | 0.003429 |
| GO:0055075 | potassium ion homeostasis | 9/3445 | 19/26032 | 0.003442 |
| GO:1905428 | regulation of plant organ formation | 9/3445 | 19/26032 | 0.003442 |
| GO:0048437 | floral organ development | 58/3445 | 281/26032 | 0.003665 |
| GO:0031408 | oxylipin biosynthetic process | 10/3445 | 23/26032 | 0.003741 |
| GO:0046470 | phosphatidylcholine metabolic process | 10/3445 | 23/26032 | 0.003741 |
| GO:0044087 | regulation of cellular component biogenesis | 36/3445 | 153/26032 | 0.00375 |
| GO:0000904 | cell morphogenesis involved in differentiation | 63/3445 | 313/26032 | 0.004193 |
| GO:0050776 | regulation of immune response | 28/3445 | 110/26032 | 0.004229 |
| GO:0048481 | plant ovule development | 18/3445 | 59/26032 | 0.004399 |
| GO:0009130 | pyrimidine nucleoside monophosphate biosynthetic process | 8/3445 | 16/26032 | 0.004515 |
| GO:0030162 | regulation of proteolysis | 21/3445 | 74/26032 | 0.004515 |
| GO:0000911 | cytokinesis by cell plate formation | 20/3445 | 69/26032 | 0.004515 |
| GO:0035556 | intracellular signal transduction | 90/3445 | 484/26032 | 0.004874 |

| | | | | |
|---|---|---|---|---|
| GO:0045944 | positive regulation of transcription by RNA polymerase II | 41/3445 | 184/26032 | 0.004951 |
| GO:0048440 | carpel development | 23/3445 | 85/26032 | 0.005121 |
| GO:0070588 | calcium ion transmembrane transport | 14/3445 | 41/26032 | 0.005121 |
| GO:0019344 | cysteine biosynthetic process | 9/3445 | 20/26032 | 0.005121 |
| GO:0005991 | trehalose metabolic process | 10/3445 | 24/26032 | 0.005274 |
| GO:0006094 | gluconeogenesis | 10/3445 | 24/26032 | 0.005274 |
| GO:0006816 | calcium ion transport | 20/3445 | 70/26032 | 0.005331 |
| GO:0010876 | lipid localization | 38/3445 | 168/26032 | 0.005553 |
| GO:0006739 | NADP metabolic process | 15/3445 | 46/26032 | 0.005571 |
| GO:0048199 | vesicle targeting, to, from or within Golgi | 7/3445 | 13/26032 | 0.005571 |
| GO:0009112 | nucleobase metabolic process | 13/3445 | 37/26032 | 0.00568 |
| GO:0009787 | regulation of abscisic acid-activated signaling pathway | 31/3445 | 129/26032 | 0.005726 |
| GO:1901419 | regulation of response to alcohol | 31/3445 | 129/26032 | 0.005726 |
| GO:1905957 | regulation of cellular response to alcohol | 31/3445 | 129/26032 | 0.005726 |
| GO:0009639 | response to red or far red light | 77/3445 | 405/26032 | 0.005755 |
| GO:1901657 | glycosyl compound metabolic process | 60/3445 | 300/26032 | 0.006025 |
| GO:0000096 | sulfur amino acid metabolic process | 18/3445 | 61/26032 | 0.006179 |
| GO:0035670 | plant-type ovary development | 18/3445 | 61/26032 | 0.006179 |
| GO:0046939 | nucleotide phosphorylation | 19/3445 | 66/26032 | 0.006195 |
| GO:0009832 | plant-type cell wall biogenesis | 63/3445 | 319/26032 | 0.00621 |
| GO:0051261 | protein depolymerization | 14/3445 | 42/26032 | 0.006216 |
| GO:0010029 | regulation of seed germination | 24/3445 | 92/26032 | 0.006382 |
| GO:0051494 | negative regulation of cytoskeleton organization | 8/3445 | 17/26032 | 0.006667 |
| GO:1902904 | negative regulation of supramolecular fiber organization | 8/3445 | 17/26032 | 0.006667 |
| GO:0006073 | cellular glucan metabolic process | 48/3445 | 229/26032 | 0.006705 |
| GO:0044042 | glucan metabolic process | 49/3445 | 235/26032 | 0.006705 |
| GO:0000910 | cytokinesis | 40/3445 | 182/26032 | 0.006788 |
| GO:0009959 | negative gravitropism | 10/3445 | 25/26032 | 0.006898 |
| GO:0031407 | oxylipin metabolic process | 10/3445 | 25/26032 | 0.006898 |
| GO:0048768 | root hair cell tip growth | 10/3445 | 25/26032 | 0.006898 |
| GO:0090630 | activation of GTPase activity | 10/3445 | 25/26032 | 0.006898 |
| GO:0032259 | methylation | 57/3445 | 284/26032 | 0.006898 |
| GO:0051017 | actin filament bundle assembly | 9/3445 | 21/26032 | 0.006903 |
| GO:0061572 | actin filament bundle organization | 9/3445 | 21/26032 | 0.006903 |
| GO:0045089 | positive regulation of innate immune response | 14/3445 | 43/26032 | 0.007625 |
| GO:0071483 | cellular response to blue light | 12/3445 | 34/26032 | 0.007823 |
| GO:0009267 | cellular response to starvation | 39/3445 | 178/26032 | 0.007988 |
| GO:0040029 | regulation of gene expression, epigenetic | 32/3445 | 138/26032 | 0.008152 |
| GO:0010608 | post-transcriptional regulation of gene expression | 54/3445 | 268/26032 | 0.008228 |
| GO:0018105 | peptidyl-serine phosphorylation | 26/3445 | 105/26032 | 0.008461 |
| GO:0009066 | aspartate family amino acid metabolic process | 18/3445 | 63/26032 | 0.008502 |
| GO:0042126 | nitrate metabolic process | 7/3445 | 14/26032 | 0.008535 |
| GO:0042128 | nitrate assimilation | 7/3445 | 14/26032 | 0.008535 |

| | | | | |
|---|---|---|---|---|
| GO:0006995 | cellular response to nitrogen starvation | 13/3445 | 39/26032 | 0.008678 |
| GO:0042546 | cell wall biogenesis | 90/3445 | 496/26032 | 0.008832 |
| GO:1900140 | regulation of seedling development | 24/3445 | 95/26032 | 0.009395 |
| GO:0009741 | response to brassinosteroid | 28/3445 | 117/26032 | 0.009477 |
| GO:0018209 | peptidyl-serine modification | 26/3445 | 106/26032 | 0.009477 |
| GO:0002682 | regulation of immune system process | 39/3445 | 180/26032 | 0.009477 |
| GO:0006206 | pyrimidine nucleobase metabolic process | 8/3445 | 18/26032 | 0.009477 |
| GO:0009065 | glutamine family amino acid catabolic process | 8/3445 | 18/26032 | 0.009477 |
| GO:0006812 | cation transport | 67/3445 | 351/26032 | 0.009565 |
| GO:0048588 | developmental cell growth | 58/3445 | 295/26032 | 0.009593 |
| GO:0032535 | regulation of cellular component size | 19/3445 | 69/26032 | 0.00961 |
| GO:0090066 | regulation of anatomical structure size | 19/3445 | 69/26032 | 0.00961 |
| GO:0043414 | macromolecule methylation | 47/3445 | 228/26032 | 0.00961 |
| GO:0048438 | floral whorl development | 46/3445 | 222/26032 | 0.00961 |
| GO:0009809 | lignin biosynthetic process | 17/3445 | 59/26032 | 0.009782 |

**Table S16.** Significantly enriched KEGG pathways of convergently contracted gene families in alpine plant genomes based on the hypergeometric test.

| **ID** | **Description** | **Gene Ratio** | **Background Ratio** | ***p*-adjust value** |
|---|---|---|---|---|
| ath00620 | Pyruvate metabolism | 39/869 | 97/5098 | 4.73E-06 |
| ath00710 | Carbon fixation in photosynthetic organisms | 30/869 | 69/5098 | 1.15E-05 |
| ath01200 | Carbon metabolism | 78/869 | 272/5098 | 2.41E-05 |
| ath00960 | Tropane, piperidine and pyridine alkaloid biosynthesis | 19/869 | 36/5098 | 2.73E-05 |
| ath01232 | Nucleotide metabolism | 31/869 | 84/5098 | 0.000197 |
| ath01230 | Biosynthesis of amino acids | 67/869 | 244/5098 | 0.000381 |
| ath00250 | Alanine, aspartate and glutamate metabolism | 21/869 | 51/5098 | 0.00053 |
| ath04075 | Plant hormone signal transduction | 76/869 | 291/5098 | 0.00053 |
| ath03015 | mRNA surveillance pathway | 38/869 | 119/5098 | 0.00053 |
| ath00270 | Cysteine and methionine metabolism | 39/869 | 124/5098 | 0.000556 |
| ath00430 | Taurine and hypotaurine metabolism | 8/869 | 11/5098 | 0.000716 |
| ath00630 | Glyoxylate and dicarboxylate metabolism | 27/869 | 78/5098 | 0.001077 |
| ath00590 | Arachidonic acid metabolism | 11/869 | 20/5098 | 0.001077 |
| ath04145 | Phagosome | 28/869 | 83/5098 | 0.001207 |
| ath00240 | Pyrimidine metabolism | 23/869 | 64/5098 | 0.001502 |
| ath04144 | Endocytosis | 44/869 | 158/5098 | 0.002732 |
| ath00010 | Glycolysis / Gluconeogenesis | 34/869 | 119/5098 | 0.007067 |
| ath00350 | Tyrosine metabolism | 15/869 | 41/5098 | 0.012192 |
| ath04136 | Autophagy - other | 15/869 | 41/5098 | 0.012192 |
| ath04120 | Ubiquitin mediated proteolysis | 40/869 | 155/5098 | 0.017982 |
| ath00564 | Glycerophospholipid metabolism | 28/869 | 99/5098 | 0.017982 |
| ath00220 | Arginine biosynthesis | 13/869 | 36/5098 | 0.023298 |
| ath03040 | Spliceosome | 47/869 | 192/5098 | 0.023298 |
| ath04626 | Plant-pathogen interaction | 49/869 | 204/5098 | 0.027774 |
| ath04016 | MAPK signaling pathway - plant | 35/869 | 139/5098 | 0.039598 |
| ath00910 | Nitrogen metabolism | 14/869 | 43/5098 | 0.040714 |
| ath00909 | Sesquiterpenoid and triterpenoid biosynthesis | 9/869 | 23/5098 | 0.040714 |

**Table S17.** Annotation information of 36 convergent selected genes. Gene functions are adopted from The Arabidopsis Information Resource (TAIR) database.

| **Orthologs** | **Species** | **Genes** | **Annotations** |
|---|---|---|---|
| OG0004760 | *C. himalaica, E. heterophyllum, H. vulgare* var. *nudum, P. mira, R. alexandrae* | *AT5G13310, AT5G13970* | Fruit development, heterocycle catabolic process |
| OG0000832 | *C. himalaica, H. vulgare* var. *nudum, P. mira, R. alexandrae, S. obvallata* | *MYB48, MYB27, MYB59* | Regulation of flavonol biosynthesis, regulation of cell cycle progression and root elongation |
| OG0012010 | *E. heterophyllum, H. vulgare* var. *nudum, P. mira, R. alexandrae, S. brachista* | *AT5G52220* | Mitotic sister chromatid cohesion |
| OG0000329 | *C. himalaica, E. heterophyllum, H. vulgare* var. *nudum, R. alexandrae* | *SEP1, SEP2, SEP3, SEP4* | Development of sepals, petals, stamens and carpels, determination of flower meristem and organ identity |
| OG0005233 | *C. himalaica, E. heterophyllum, R. alexandrae, S. brachista* | *BBX11* | Circadian rhythm, response to blue light, response to light stimulus, response to red light |
| OG0000556 | *C. himalaica, H. vulgare* var. *nudum, P. mira, S. brachista* | *AT5G21090, AT3G43740* | Signal transduction |
| OG0001948 | *C. himalaica, H. vulgare* var. *nudum, R. alexandrae, S. brachista* | *AT5P1, AT1G71710* | Phosphatidylinositol dephosphorylation, seed germination, seedling development |
| OG0001928 | *C. himalaica, P. mira, S. brachista, S. obvallata* | *PMDH1, PMDH2* | Regulation of fatty acid beta-oxidation, regulation of photorespiration |
| OG0009364 | *E. heterophyllum, H. vulgare* var. *nudum, P. mira, R. alexandrae* | *OHP2* | Photosystem II assembly, response to light intensity |
| OG0009704 | *H. vulgare* var. *nudum, P. mira, R. alexandrae, S. obvallata* | *HSA32, AT4G21323* | Heat acclimation, response to heat |
| OG0007885 | *C. himalaica, E. heterophyllum, H. vulgare* var. *nudum* | *SVL4, SVL5* | Pollen tube growth, lipid metabolic process |
| OG0005129 | *C. himalaica, E. heterophyllum, H. vulgare* var. *nudum* | *AT3G55470* | Regulation of defense response, response to water deprivation, shoot system development |
| OG0004460 | *C. himalaica, E. heterophyllum, P. mira* | *LIN37A, LIN37B* | Cellular component biogenesis, organelle organization |
| OG0004639 | *C. himalaica, E. heterophyllum, S. brachista* | *EMB2777, AT3G28230* | Maturation of SSU-rrna from tricistronic rrna transcript (SSU-rrna, 5.8S rrna, LSU-rrna) |
| OG0001094 | *C. himalaica, H. vulgare* var. *nudum, S. brachista* | *NUC-L1, NUC-L2* | Rrna processing, leaf development, root development, shoot system development |
| OG0006408 | *C. himalaica, H. vulgare* var. *nudum, S. brachista* | *AT1G12730, AT1G63110* | Response to temperature stimulus, protein localization to cell surface |

| | | | |
|---|---|---|---|
| OG0001067 | *C. himalaica, H. vulgare* var. *nudum, S. brachista* | *CAT1, CAT2, CAT3* | Response to cold, response to light stimulus, response to hydrogen peroxide, response to oxidative stress |
| OG0007544 | *C. himalaica, P. mira, S. brachista* | *LSP1* | Response to virus, translational initiation |
| OG0006996 | *C. himalaica, P. mira, S. brachista* | *HPR* | Photorespiration, cellular response to light stimulus |
| OG0000803 | *C. himalaica, P. mira, S. obvallata* | *AEP1, AEP2, AEP3* | Leaf senescence, protein catabolic process, response to ethylene |
| OG0001859 | *C. himalaica, R. alexandrae, S. obvallata* | *AT1G32610, VQ9, IKU1, AT5G46780* | Endosperm development, regulation of seed growth |
| OG0001302 | *C. himalaica, R. alexandrae, S. obvallata* | *KPNB1, AT3G08947, AT3G08943* | Protein import into nucleus, drought stress |
| OG0003962 | *E. heterophyllum, S. brachista, S. obvallata* | *DUF726, AT4G36210* | Response to lipid, shoot system development, signal transduction |
| OG0002378 | *H. vulgare* var. *nudum, P. mira, S. brachista* | *SLP3, AT4G30020* | Cell differentiation, cell division, proteolysis, tissue development |
| OG0007327 | *H. vulgare* var. *nudum, P. mira, S. brachista* | *TPC1* | Calcium ion transport, regulation of stomatal movement, seed germination |
| OG0000811 | *H. vulgare* var. *nudum, R. alexandrae, S. brachista* | *NOT4A, NOT4B, NOT4C* | Protein ubiquitination, regulation of chloroplast protein synthesis |
| OG0001877 | *H. vulgare* var. *nudum, R. alexandrae, S. brachista* | *PRP4KA, PRP4KB, AT3G53640* | Mrna cis splicing, via spliceosome, protein phosphorylation |
| OG0012347 | *H. vulgare* var. *nudum, R. alexandrae, S. brachista* | *AT5G64730* | Regulation of developmental process, mrna splicing, via spliceosome |
| OG0001378 | *H. vulgare* var. *nudum, S. brachista, S. obvallata* | *AT1G49750, AT3G19320, AT4G06744* | Response to jasmonic acid, response to wounding, secondary metabolic process |
| OG0003532 | *H. vulgare* var. *nudum, S. brachista, S. obvallata* | *MMD1, AT2G01810, AT2G07714, ATMG00550* | Meiotic cell cycle, male meiosis chromosome segregation, pollen sperm cell differentiation |
| OG0005907 | *H. vulgare* var. *nudum, S. brachista, S. obvallata* | *AT3G06868, AT5G49100* | Cell wall biogenesis, plant-type cell wall organization or biogenesis |
| OG0001368 | *P. mira, R. alexandrae, S. brachista* | *AT1G14670, AT2G01970, AT5G37310* | Protein localization to membrane |
| OG0004363 | *P. mira, R. alexandrae, S. obvallata* | *PEPKR2* | Intracellular signal transduction, peptidyl-serine phosphorylation, protein autophosphorylation |
| OG0001613 | *P. mira, S. brachista, S. obvallata* | *PME21, PMEI-PME28, PME58, AT3G06830* | Cell wall modification, pectin catabolic process, seed coat development |
| OG0006829 | *P. mira, S. brachista, S. obvallata* | *AT5G14580* | RNA catabolic process, mitochondrial RNA processing |
| OG0004220 | *R. alexandrae, S. brachista, S. obvallata* | *NBS1* | DNA repair, cellular response to DNA damage stimulus, double-strand break repair via homologous recombination, mitotic recombination |

**Table S18.** Significantly enriched GO terms of genes undergoing convergent positive selection in alpine plant genomes.

| ID | Description | Gene Ratio | Background Ratio | *p*-adjust value |
|---|---|---|---|---|
| GO:0009970 | cellular response to sulfate starvation | 3/75 | 14/26032 | 0.003228 |
| GO:0019217 | regulation of fatty acid metabolic process | 3/75 | 28/26032 | 0.007984 |
| GO:0045292 | mRNA cis splicing, via spliceosome | 3/75 | 31/26032 | 0.007984 |
| GO:0042744 | hydrogen peroxide catabolic process | 3/75 | 32/26032 | 0.007984 |
| GO:0048440 | carpel development | 4/75 | 85/26032 | 0.007984 |
| GO:0046488 | phosphatidylinositol metabolic process | 4/75 | 91/26032 | 0.007984 |
| GO:0030258 | lipid modification | 4/75 | 95/26032 | 0.007984 |
| GO:0045490 | pectin catabolic process | 4/75 | 98/26032 | 0.007984 |
| GO:0006995 | cellular response to nitrogen starvation | 3/75 | 39/26032 | 0.007984 |
| GO:0048467 | gynoecium development | 4/75 | 100/26032 | 0.007984 |
| GO:0009853 | photorespiration | 3/75 | 46/26032 | 0.011032 |
| GO:0043562 | cellular response to nitrogen levels | 3/75 | 48/26032 | 0.011032 |
| GO:0050994 | regulation of lipid catabolic process | 2/75 | 10/26032 | 0.011032 |
| GO:0006606 | protein import into nucleus | 3/75 | 55/26032 | 0.014248 |
| GO:0042743 | hydrogen peroxide metabolic process | 3/75 | 55/26032 | 0.014248 |
| GO:0051170 | import into nucleus | 3/75 | 58/26032 | 0.015241 |
| GO:0048481 | plant ovule development | 3/75 | 59/26032 | 0.015241 |
| GO:0016485 | protein processing | 3/75 | 60/26032 | 0.015241 |
| GO:0035670 | plant-type ovary development | 3/75 | 61/26032 | 0.015241 |
| GO:0000272 | polysaccharide catabolic process | 4/75 | 143/26032 | 0.015489 |
| GO:0080113 | regulation of seed growth | 2/75 | 15/26032 | 0.015787 |
| GO:0034504 | protein localization to nucleus | 3/75 | 68/26032 | 0.018067 |
| GO:0006650 | glycerophospholipid metabolic process | 4/75 | 159/26032 | 0.019216 |
| GO:0006108 | malate metabolic process | 2/75 | 18/26032 | 0.019216 |
| GO:0080112 | seed growth | 2/75 | 18/26032 | 0.019216 |
| GO:0042542 | response to hydrogen peroxide | 3/75 | 76/26032 | 0.021093 |
| GO:0045488 | pectin metabolic process | 4/75 | 171/26032 | 0.021603 |
| GO:0010393 | galacturonan metabolic process | 4/75 | 172/26032 | 0.021603 |
| GO:0042545 | cell wall modification | 4/75 | 173/26032 | 0.021603 |
| GO:0043094 | cellular metabolic compound salvage | 3/75 | 81/26032 | 0.021959 |
| GO:0016036 | cellular response to phosphate starvation | 3/75 | 82/26032 | 0.022013 |
| GO:0045944 | positive regulation of transcription by RNA polymerase II | 4/75 | 184/26032 | 0.023147 |
| GO:0046855 | inositol phosphate dephosphorylation | 2/75 | 23/26032 | 0.023147 |
| GO:0071545 | inositol phosphate catabolic process | 2/75 | 23/26032 | 0.023147 |
| GO:0071555 | cell wall organization | 5/75 | 313/26032 | 0.023848 |
| GO:0019216 | regulation of lipid metabolic process | 3/75 | 90/26032 | 0.024735 |
| GO:0046486 | glycerolipid metabolic process | 4/75 | 197/26032 | 0.026456 |
| GO:0046838 | phosphorylated carbohydrate | 2/75 | 26/26032 | 0.026456 |

| | | | | |
|---|---|---|---|---|
| | dephosphorylation | | | |
| GO:0051604 | protein maturation | 3/75 | 96/26032 | 0.027094 |
| GO:0046856 | phosphatidylinositol dephosphorylation | 2/75 | 27/26032 | 0.027094 |
| GO:0006506 | GPI anchor biosynthetic process | 2/75 | 30/26032 | 0.031801 |
| GO:0046839 | phospholipid dephosphorylation | 2/75 | 30/26032 | 0.031801 |
| GO:0006505 | GPI anchor metabolic process | 2/75 | 31/26032 | 0.033142 |
| GO:0048438 | floral whorl development | 4/75 | 222/26032 | 0.035007 |
| GO:0046174 | polyol catabolic process | 2/75 | 33/26032 | 0.035826 |
| GO:0006644 | phospholipid metabolic process | 4/75 | 228/26032 | 0.036311 |
| GO:0045229 | external encapsulating structure organization | 5/75 | 370/26032 | 0.036311 |
| GO:0016052 | carbohydrate catabolic process | 4/75 | 230/26032 | 0.036374 |
| GO:0010565 | regulation of cellular ketone metabolic process | 3/75 | 117/26032 | 0.038068 |
| GO:0006913 | nucleocytoplasmic transport | 3/75 | 121/26032 | 0.040125 |
| GO:0051169 | nuclear transport | 3/75 | 121/26032 | 0.040125 |
| GO:0031668 | cellular response to extracellular stimulus | 4/75 | 243/26032 | 0.040125 |
| GO:0000462 | maturation of SSU-rRNA from tricistronic rRNA transcript (SSU-rRNA, 5.8S rRNA, LSU-rRNA) | 2/75 | 38/26032 | 0.040125 |
| GO:0000398 | mRNA splicing, via spliceosome | 4/75 | 246/26032 | 0.040754 |
| GO:0098813 | nuclear chromosome segregation | 3/75 | 125/26032 | 0.040754 |
| GO:0071496 | cellular response to external stimulus | 4/75 | 249/26032 | 0.041196 |
| GO:0043647 | inositol phosphate metabolic process | 2/75 | 40/26032 | 0.041243 |
| GO:0044282 | small molecule catabolic process | 4/75 | 253/26032 | 0.041699 |
| GO:0046164 | alcohol catabolic process | 2/75 | 41/26032 | 0.041699 |
| GO:0072593 | reactive oxygen species metabolic process | 3/75 | 131/26032 | 0.041699 |
| GO:0006635 | fatty acid beta-oxidation | 2/75 | 42/26032 | 0.041699 |
| GO:0009960 | endosperm development | 2/75 | 42/26032 | 0.041699 |
| GO:0007059 | chromosome segregation | 3/75 | 137/26032 | 0.045802 |
| GO:0000375 | RNA splicing, via transesterification reactions | 4/75 | 272/26032 | 0.048201 |
| GO:0000377 | RNA splicing, via transesterification reactions with bulged adenosine as nucleophile | 4/75 | 272/26032 | 0.048201 |

**Table S19.** Significantly enriched KEGG pathways of genes undergoing convergent positive selection in alpine plant genomes.

| ID | Description | Gene Ratio | Background Ratio | *p*-adjust value |
|---|---|---|---|---|
| ath00630 | Glyoxylate and dicarboxylate metabolism | 6/21 | 78/5105 | 7.58E-06 |
| ath01200 | Carbon metabolism | 6/21 | 272/5105 | 0.004073 |
| ath00040 | Pentose and glucuronate interconversions | 4/21 | 108/5105 | 0.004073 |
| ath03018 | RNA degradation | 4/21 | 113/5105 | 0.004073 |
| ath00380 | Tryptophan metabolism | 3/21 | 64/5105 | 0.006808 |
| ath00563 | Glycosylphosphatidylinositol (GPI)-anchor biosynthesis | 2/21 | 26/5105 | 0.011642 |
| ath04146 | Peroxisome | 3/21 | 87/5105 | 0.011642 |
| ath03013 | Nucleocytoplasmic transport | 3/21 | 103/5105 | 0.016293 |
| ath04016 | MAPK signaling pathway - plant | 3/21 | 139/5105 | 0.032616 |
| ath00020 | Citrate cycle (TCA cycle) | 2/21 | 64/5105 | 0.044605 |
| ath00710 | Carbon fixation in photosynthetic organisms | 2/21 | 69/5105 | 0.046611 |

**Table S20.** Significantly enriched GO terms of gene families undergoing convergently accelerated evolutionary rate in alpine plant genomes.

| **ID** | **Description** | **Gene Ratio** | **Background Ratio** | ***p*-adjust value** |
|---|---|---|---|---|
| GO:0048544 | recognition of pollen | 27/255 | 50/26032 | 6.72E-39 |
| GO:0008037 | cell recognition | 27/255 | 52/26032 | 1.46E-38 |
| GO:0009875 | pollen-pistil interaction | 27/255 | 72/26032 | 8.09E-34 |
| GO:0009856 | pollination | 35/255 | 374/26032 | 5.89E-22 |
| GO:0044706 | multi-multicellular organism process | 35/255 | 374/26032 | 5.89E-22 |
| GO:0009696 | salicylic acid metabolic process | 11/255 | 52/26032 | 2.46E-10 |
| GO:0006955 | immune response | 20/255 | 283/26032 | 5.40E-10 |
| GO:0009694 | jasmonic acid metabolic process | 11/255 | 57/26032 | 5.40E-10 |
| GO:0045087 | innate immune response | 18/255 | 237/26032 | 1.52E-09 |
| GO:0018958 | phenol-containing compound metabolic process | 11/255 | 69/26032 | 3.85E-09 |
| GO:0042537 | benzene-containing compound metabolic process | 11/255 | 76/26032 | 1.04E-08 |
| GO:0009736 | cytokinin-activated signaling pathway | 10/255 | 80/26032 | 2.78E-07 |
| GO:0071368 | cellular response to cytokinin stimulus | 10/255 | 86/26032 | 5.25E-07 |
| GO:0009735 | response to cytokinin | 10/255 | 133/26032 | 3.11E-05 |
| GO:0032270 | positive regulation of cellular protein metabolic process | 10/255 | 138/26032 | 4.07E-05 |
| GO:0051247 | positive regulation of protein metabolic process | 10/255 | 149/26032 | 7.67E-05 |
| GO:0031408 | oxylipin biosynthetic process | 5/255 | 23/26032 | 8.07E-05 |
| GO:0031407 | oxylipin metabolic process | 5/255 | 25/26032 | 0.000118 |
| GO:0032268 | regulation of cellular protein metabolic process | 15/255 | 393/26032 | 0.00025 |
| GO:0070534 | protein K63-linked ubiquitination | 4/255 | 15/26032 | 0.000306 |
| GO:0051246 | regulation of protein metabolic process | 15/255 | 419/26032 | 0.000457 |
| GO:0031398 | positive regulation of protein ubiquitination | 4/255 | 17/26032 | 0.000457 |
| GO:1903322 | positive regulation of protein modification by small protein conjugation or removal | 4/255 | 17/26032 | 0.000457 |
| GO:0006301 | postreplication repair | 4/255 | 18/26032 | 0.000559 |
| GO:0044772 | mitotic cell cycle phase transition | 7/255 | 89/26032 | 0.000602 |
| GO:1901615 | organic hydroxy compound metabolic process | 15/255 | 440/26032 | 0.000681 |
| GO:0072350 | tricarboxylic acid metabolic process | 4/255 | 21/26032 | 0.000949 |
| GO:0003008 | system process | 4/255 | 22/26032 | 0.000971 |
| GO:0003013 | circulatory system process | 4/255 | 22/26032 | 0.000971 |
| GO:0003018 | vascular process in circulatory system | 4/255 | 22/26032 | 0.000971 |
| GO:0010232 | vascular transport | 4/255 | 22/26032 | 0.000971 |
| GO:0010233 | phloem transport | 4/255 | 22/26032 | 0.000971 |
| GO:0031396 | regulation of protein ubiquitination | 4/255 | 23/26032 | 0.001132 |
| GO:1903320 | regulation of protein modification by small protein conjugation or removal | 4/255 | 24/26032 | 0.001308 |
| GO:0035556 | intracellular signal transduction | 15/255 | 484/26032 | 0.001459 |
| GO:0090630 | activation of GTPase activity | 4/255 | 25/26032 | 0.001459 |
| GO:0000209 | protein polyubiquitination | 7/255 | 111/26032 | 0.001671 |

| | | | | |
|---|---|---|---|---|
| GO:0044770 | cell cycle phase transition | 7/255 | 112/26032 | 0.001721 |
| GO:0031399 | regulation of protein modification process | 9/255 | 197/26032 | 0.002006 |
| GO:0007623 | circadian rhythm | 8/255 | 155/26032 | 0.002006 |
| GO:0048511 | rhythmic process | 8/255 | 155/26032 | 0.002006 |
| GO:0000079 | regulation of cyclin-dependent protein serine/threonine kinase activity | 5/255 | 54/26032 | 0.002341 |
| GO:1904029 | regulation of cyclin-dependent protein kinase activity | 5/255 | 54/26032 | 0.002341 |
| GO:0071900 | regulation of protein serine/threonine kinase activity | 5/255 | 59/26032 | 0.00348 |
| GO:0000160 | phosphorelay signal transduction system | 7/255 | 130/26032 | 0.003653 |
| GO:0006448 | regulation of translational elongation | 3/255 | 14/26032 | 0.003685 |
| GO:0010114 | response to red light | 6/255 | 101/26032 | 0.00561 |
| GO:0043547 | positive regulation of GTPase activity | 4/255 | 40/26032 | 0.007046 |
| GO:0009309 | amine biosynthetic process | 4/255 | 42/26032 | 0.008159 |
| GO:0042401 | cellular biogenic amine biosynthetic process | 4/255 | 42/26032 | 0.008159 |
| GO:0045859 | regulation of protein kinase activity | 5/255 | 74/26032 | 0.008578 |
| GO:0043087 | regulation of GTPase activity | 4/255 | 43/26032 | 0.008584 |
| GO:0043549 | regulation of kinase activity | 5/255 | 76/26032 | 0.009317 |
| GO:0051345 | positive regulation of hydrolase activity | 4/255 | 46/26032 | 0.010682 |
| GO:0006415 | translational termination | 3/255 | 21/26032 | 0.010704 |
| GO:0043244 | regulation of protein-containing complex disassembly | 3/255 | 24/26032 | 0.015656 |
| GO:0045727 | positive regulation of translation | 3/255 | 25/26032 | 0.017353 |
| GO:0001932 | regulation of protein phosphorylation | 5/255 | 90/26032 | 0.018133 |
| GO:0034250 | positive regulation of cellular amide metabolic process | 3/255 | 27/26032 | 0.021016 |
| GO:0042325 | regulation of phosphorylation | 5/255 | 97/26032 | 0.024323 |
| GO:0042752 | regulation of circadian rhythm | 4/255 | 61/26032 | 0.026852 |
| GO:0050790 | regulation of catalytic activity | 9/255 | 302/26032 | 0.026852 |
| GO:0051726 | regulation of cell cycle | 8/255 | 247/26032 | 0.026852 |
| GO:0051338 | regulation of transferase activity | 5/255 | 101/26032 | 0.026852 |
| GO:0031401 | positive regulation of protein modification process | 4/255 | 62/26032 | 0.026852 |
| GO:0016126 | sterol biosynthetic process | 3/255 | 33/26032 | 0.033097 |
| GO:1990573 | potassium ion import across plasma membrane | 2/255 | 10/26032 | 0.033097 |
| GO:0006545 | glycine biosynthetic process | 2/255 | 11/26032 | 0.03366 |
| GO:0007094 | mitotic spindle assembly checkpoint signaling | 2/255 | 11/26032 | 0.03366 |
| GO:0031577 | spindle checkpoint signaling | 2/255 | 11/26032 | 0.03366 |
| GO:0033046 | negative regulation of sister chromatid segregation | 2/255 | 11/26032 | 0.03366 |
| GO:0033047 | regulation of mitotic sister chromatid segregation | 2/255 | 11/26032 | 0.03366 |
| GO:0033048 | negative regulation of mitotic sister chromatid segregation | 2/255 | 11/26032 | 0.03366 |
| GO:0045841 | negative regulation of mitotic metaphase/anaphase transition | 2/255 | 11/26032 | 0.03366 |
| GO:0051985 | negative regulation of chromosome segregation | 2/255 | 11/26032 | 0.03366 |
| GO:0071173 | spindle assembly checkpoint signaling | 2/255 | 11/26032 | 0.03366 |
| GO:0071174 | mitotic spindle checkpoint signaling | 2/255 | 11/26032 | 0.03366 |
| GO:1902100 | negative regulation of metaphase/anaphase transition of cell cycle | 2/255 | 11/26032 | 0.03366 |
| GO:1905819 | negative regulation of chromosome separation | 2/255 | 11/26032 | 0.03366 |
| GO:2000816 | negative regulation of mitotic sister chromatid separation | 2/255 | 11/26032 | 0.03366 |

| | | | | |
|---|---|---|---|---|
| GO:0012501 | programmed cell death | 6/255 | 161/26032 | 0.034837 |
| GO:0019430 | removal of superoxide radicals | 2/255 | 12/26032 | 0.037772 |
| GO:0071450 | cellular response to oxygen radical | 2/255 | 12/26032 | 0.037772 |
| GO:0071451 | cellular response to superoxide | 2/255 | 12/26032 | 0.037772 |
| GO:0071484 | cellular response to light intensity | 2/255 | 12/26032 | 0.037772 |
| GO:0048278 | vesicle docking | 3/255 | 38/26032 | 0.038421 |
| GO:0065009 | regulation of molecular function | 9/255 | 338/26032 | 0.039785 |
| GO:0008219 | cell death | 7/255 | 231/26032 | 0.048022 |
| GO:0000303 | response to superoxide | 2/255 | 14/26032 | 0.048022 |
| GO:0000305 | response to oxygen radical | 2/255 | 14/26032 | 0.048022 |
| GO:2001251 | negative regulation of chromosome organization | 2/255 | 14/26032 | 0.048022 |
| GO:0046777 | protein autophosphorylation | 6/255 | 177/26032 | 0.048048 |
| GO:0071478 | cellular response to radiation | 6/255 | 178/26032 | 0.048798 |

**Table S21.** Significantly enriched KEGG pathways of gene families undergoing convergently accelerated evolutionary rate in alpine plant genomes.

| **ID** | **Description** | **Gene Ratio** | **Background Ratio** | ***p*-adjust value** |
|---|---|---|---|---|
| ath00040 | Pentose and glucuronate interconversions | 12/57 | 107/5095 | 3.05E-08 |
| ath03430 | Mismatch repair | 5/57 | 37/5095 | 0.000584 |
| ath03030 | DNA replication | 5/57 | 48/5095 | 0.001392 |
| ath04075 | Plant hormone signal transduction | 11/57 | 289/5095 | 0.001717 |
| ath03440 | Homologous recombination | 5/57 | 64/5095 | 0.003245 |
| ath03420 | Nucleotide excision repair | 5/57 | 67/5095 | 0.003339 |
| ath04144 | Endocytosis | 7/57 | 158/5095 | 0.005942 |
| ath00999 | Biosynthesis of various plant secondary metabolites | 4/57 | 64/5095 | 0.016304 |
| ath00073 | Cutin, suberine and wax biosynthesis | 3/57 | 37/5095 | 0.021013 |
| ath00940 | Phenylpropanoid biosynthesis | 5/57 | 128/5095 | 0.032743 |
| ath04130 | SNARE interactions in vesicular transport | 3/57 | 47/5095 | 0.033166 |
| ath00670 | One carbon pool by folate | 2/57 | 21/5095 | 0.045072 |

**Table S22.** Tests for relaxed selection on *S*-locus genes of alpine plants using RELAX model. *k*: estimated selection intensity. *p*-values are calculated by likelihood ratio test and significant results ($p < 0.05$) are in bold.

| **Species** | ***k*** | ***p*-value** |
|---|---|---|
| ***C. himalaica*** | **0.78** | **0.025** |
| ***E. heterophyllum*** | **0.43** | **0.042** |
| ***H. vulgare* var. *nudum*** | **0.15** | **0.002** |
| *P. mira* | 1.54 | 0.078 |
| *R. alexandrae* | 0.80 | 0.201 |
| ***S. brachista*** | **0.60** | **0.001** |
| ***S. obvallata*** | **0.72** | **0.007** |

**Table S23.** Functional annotation of genes predicted to interact with *MYB48*.

| Gene | Identifier | Annotations |
|---|---|---|
| *DFR* | AT5G42800.1 | Bifunctional dihydroflavonol 4-reductase/flavanone 4-reductase; Dihydroflavonol reductase. Catalyzes the conversion of dihydroquercetin to leucocyanidin in the biosynthesis of anthocyanins |
| *F3H* | AT3G51240.1 | Naringenin,2-oxoglutarate 3-dioxygenase; Catalyzes the 3-beta-hydroxylation of 2S-flavanones to 2R,3R-dihydroflavonols which are intermediates in the biosynthesis of flavonols, anthocyanidins, catechins and proanthocyanidins in plants |
| *FLS1* | AT5G08640.1 | Flavonol synthase/flavanone 3-hydroxylase; Catalyzes the formation of flavonols from dihydroflavonols. It can act on dihydrokaempferol to produce kaempferol, on dihydroquercetin to produce quercitin and on dihydromyricetin to produce myricetin. |
| *PEI1* | AT5G07500.1 | Zinc finger C-x8-C-x5-C-x3-H type family protein; Embryo-specific transcription factor required at the globular to heart stage transition in embryo development |
| *TT5* | AT3G55120.1 | Chalcone-flavanone isomerase family protein; Catalyzes the intramolecular cyclization of bicyclic chalcones into tricyclic (S)-flavanones. Responsible for the isomerization of 4,2',4',6'-tetrahydroxychalcone (also termed chalcone) into naringenin |
| *TTG1* | AT5G24520.1 | Transducin/WD40 repeat-like superfamily protein; Required for the accumulation of purple anthocyanins in leaves and stems. Involved in trichome and root hair development. Controls epidermal cell fate specification. Affects dihydroflavonol 4-reductase gene expression. |
| *TTG2* | AT2G37260.1 | Wrky family transcription factor family protein; Encodes a protein similar to WRKY transcription factors that is expressed in the seed integument and endosperm. Mutants are defective in proanthocyanidin synthesis and seed mucilate deposition. Seeds are yellow colored |
| *AT1G15810* | AT1G15810.1 | S15/NS1, RNA-binding protein; Its function is described as structural constituent of ribosome; Involved in translation |
| *AT1G80620* | AT1G80620.1 | Uncharacterized protein T21F11.5; S15/NS1, RNA-binding protein; Its function is described as structural constituent of ribosome; Involved in translation |
| *AT3G06880* | AT3G06880.2 | Transducin/WD40 repeat-like superfamily protein; Its function is described as binding, nucleotide binding; Located in cellular |
